# Functional and structural characterization of dendritic spine pathology in a mouse model of tauopathy

**DOI:** 10.64898/2026.06.08.730960

**Authors:** Liam M. Adsit, Kyle Cekada, Ikuko T. Smith

**Affiliations:** Department of Molecular, Cellular & Developmental Biology, University of California, Santa Barbara, Santa Barbara CA 93106; Neuroscience Research Institute, University of California, Santa Barbara, Santa Barbara CA 93106; Department of Psychological & Brain Sciences, University of California, Santa Barbara, Santa Barbara CA 93106

**Keywords:** tauopathy, dendrite, dendritic spine, spine loss, spine pathology, input mapping

## Abstract

Abnormal deposition of the microtubule-associated protein tau has long been associated with neurodegenerative diseases. While spine loss and neuronal death are hallmarks of tauopathy, how pathological tau affects synaptic activity *in vivo* and whether functional properties of individual synapses dictate the survival fate of dendritic spines remain elusive. Here we examined the visual response properties of dendrites and spines of layer 2/3 primary visual cortical neurons, using longitudinal two-photon calcium imaging in P301S mouse model of tauopathy. We found that neuronal outputs in tau mutant mice were hyperactive and poorly tuned whereas dendritic spine responses were also poorly tuned but hypoactive. Moreover, we found that spines that were stably retained across two imaging sessions were larger in size and more sharply tuned but less active compared to those that turned over in controls. Such structure-to-function relationship was not observed in mutants. Our findings illustrate how the preferential maintenance of well-tuned inputs in healthy neural circuitry may be affected by tauopathy, resulting in neurons with poorly tuned visual responses.

## INTRODUCTION

Tauopathies are neurodegenerative diseases characterized by neurofibrillary tangles (NFTs) of the microtubule-associated protein tau (MAPT; reviewed in^1–4^). Perhaps best known as one of the two cardinal features of Alzheimer’s disease (AD) with the other being amyloid-beta (Aβ) plaques^5–9^, filamentous tau inclusion has been reported in a variety of brain disorders, including frontotemporal dementia with Parkinsonism linked to chromosome17 (FTDP-17), Dementia with Lewy Bodies (DLB), Parkinson’s disease (PD), and Huntington’s disease (HD) as well as in traumatic brain injury (TBI), chronic traumatic encephalopathy (CTE), depression, and chronic substance abuse^10–18^. Thus, whether of primary causes (e.g. mutations in the MAPT gene in FTD) or secondary causes (e.g. following a wide range of inflammatory triggers as in CTE), tauopathy emerges as a key mechanistic player shared among these seemingly diverse disorders^1,3,4,19^. Of note, recent studies have identified soluble tau, rather than the NFTs, as the culprit for mediating neurotoxicity and the disruption of neural functions, advancing the predicted timing of the pathological onset (reviewed in^8,9,20^). In a healthy neuron, soluble tau is more concentrated in axons than in somatodendritic compartments. By contrast in tauopathy, tau is mislocalized to dendritic spines and disrupts synaptic function^21^. *In vitro* studies have identified both presynaptic and postsynaptic origins of synapse dysfunction, including structural loss of afferent inputs, alterations in the presynaptic neurotransmitter release and reuptake, pathological activation of glutamate receptors, and calcium overload in the postsynaptic neuron^22,23^. Thus, dendritic localization of abnormal tau is directly linked to the dendritic spine pathogenesis^21^.

Visual impairments are a common symptom observed in many neurodegenerative diseases with concomitant tauopathy, including AD, FTD, DLB, PD, HD, TBI and CTE. While some of the visual dysfunction stems from the combination of progressive loss of cells in the retina and compromised optic nerves^24,25^, visual impairments due to cortical degeneration are evident in many, including contrast sensitivity^26^, loss of motion vision, poor performance in visuo-spatial tasks, and visual hallucinations^27–32^. As such, early presymptomatic onset of visual impairments has garnered attention as potential predictors of preclinical neurodegenerative disease and cognitive decline^33–35^. Using P301S mutant human tau expressing transgenic mice (PS19 line) as a model of tauopathy, we investigated the impact of pathological tau on visual response properties of layer 2/3 cortical neurons in the primary visual cortex (V1). Unlike other models of tauopathy, the optic nerve in P301S mutants is only mildly impacted by the pathological tau, and the mouse strain is spared from retinal degeneration, making them an ideal model to study the brain-specific impact of tau^36^. Of the over 30 tau gene mutations deemed pathogenic for FTDP-17, PS19 line harbors the P301S mutation in human tau driven by the mouse prion promoter, allowing for ubiquitous expression of the pathological tau across the brain including the hippocampus, amygdala, and the neocortex^37^. The P301S transgene encodes four microtubule-binding domains and one N-terminal insert (4R/1N) and contains a missense mutation in exon 10 within the 2R region, which renders the mutant tau to be susceptible to hyperphosphorylation. The signature phenotype of P301S includes accelerated onset of gross forebrain atrophy, dendritic spine loss and neuronal degeneration, leading to premature death with an average lifespan of 11-15 months^37^. Molecularly, phosphorylated tau is detected, displaced in somatodendritic compartments, a few months prior to the aggregation of NFTs ^38,39^. While areas like the hippocampus and the entorhinal cortex are affected early, resulting in extensive neurodegeneration and atrophy by 9 months of age, visual cortex exhibits modest accumulation of hyperphosphorylated tau by 6 months^37,40,41^, which progressively becomes insoluble by 12 months when significant cortical degeneration is observed^37,40^. The slow progressive spread of the pathological tau reported in the visual cortex makes P301S an ideal model for studying the early onset of visual impairment in tauopathy.

Here, using two-photon calcium (2p-Ca^2+^) imaging of dendritic shaft and spine activity with a genetically encoded calcium indicator (GECI), we provide both structural and functional characterization of the dendritic spine pathology *in vivo* in the mouse tauopathy model. Specifically, we sought to determine whether there is a link between the functional properties of the dendritic spine and their turnover, and how the spine dysfunction may affect the neuronal output.

## RESULTS

### P301S mutant model of tauopathy exhibits hyperphosphorylated tau in the primary visual cortex

We first examined the level of pathological tau expression in this mouse model of tauopathy. As expected, immunohistochemical labeling with AT8 antibodies targeting pSer202/pThr205 tau^42^ revealed extensive expression of hyperphosphorylated tau aggregates in the hippocampal formation, including the hippocampus, dentate gyrus, subiculum and entorhinal cortex in the P301S mutant human tau transgenic heterozygotes (PS19 line; **Figure s1a**). We also observed more modest but significant amount of hyperphosphorylated tau expressed in the visual cortex, including V1 (**Figure 1a**). Immunoreactivity against AT8 antibodies was found in abundance in cortical layers 2/3 and 5 (**Figure 1b, Figure S1b, c**). High-magnification images of the layer 2/3 revealed strong perisomatic expression of hyperphosphorylated tau throughout the age range used in the study that was not observed in the wildtype control mice (P301S controls, two cortical sections each from 6 mice v.s. P301S mutants, two cortical sections each from 7 mice. AT8 fluorescence intensity (a.u.) mean ± s.e.m.: controls 51.299 ± 3.994 vs. mutants 77.796 ± 7.443, two-tailed Wilcoxon test, *p =* 0.0015; **Figure 1c & d**). Phosphorylated tau expression was also observed in the dendrites of layer 2/3 neurons in mutants using STED super-resolution microscopy (**Figure 1e & f, Figure S1d & e**) revealing dendritic structures consistent with putative spines. In addition, phosphorylated tau was detected in the long apical trunks of layer 5 neurons imaged with confocal microscopy (**Figure S1b**). Despite the aberrant aggregation of hyperphosphorylated tau in the somatodendritic compartments, the anatomical structure of the cortex was grossly intact with no signs of early neural degeneration observed in the mutants (**Figure 1a – c, Figure S1a & b**). These observations confirm the validity of the use of P301S mice for investigating the direct functional impact of pathological tau in visual processing, preceding notable neuronal death.

**Figure 1.**
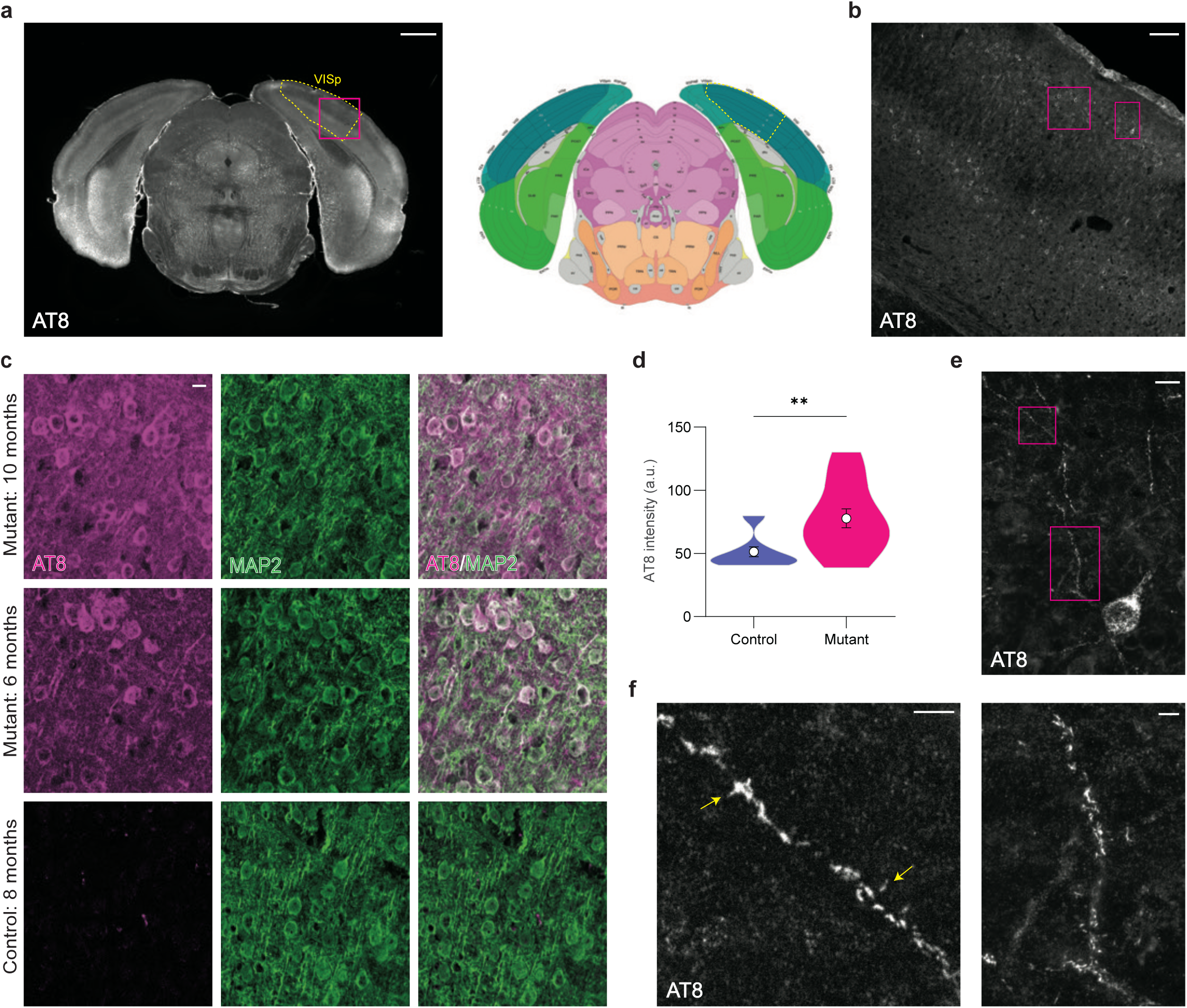
Pathological tau expression in the visual cortex of P301S mutants. (A) Left: Example coronal slice from a 10-month P301S mutant showing phosphorylated tau labeled with AT8 antibodies. Dotted yellow line outlines V1 (VISp) of right hemisphere. Scale bar = 1 mm. Right: A slice from Allen Mouse Brain Reference Atlas (ref.75) corresponding to the equivalent coordinates shown for an anatomical reference. See also Figure S1a. (B) Magnified view of magenta box from panel (A), showing phosphorylated tau expression across layers 1-6 of V1. Scale bar = 100 µm. See also Figure S1b & c. (C) Example images from V1 layer 2/3 neurons from control (8 months) and mutant mice (6 and 10 months). The images show the overlap of phosphorylated tau with MAP2, a neuronal marker used to reveal cytoarchitecture. The top left image is a magnified view of left magenta box from panel (B). Scale bar = 10 µm. (D) Quantification of AT8 fluorescence intensity in V1 layer 2/3 in control and mutant mice. Mutants (n = 14 slices) had significantly higher AT8 intensity compared to controls (n = 12 slices). (E) Magnified view of right magenta box from panel (B) acquired with STED microscopy, showing an example V1 layer 2/3 neuron with robust phosphorylated tau expression in soma and dendrites. Scale bar = 2 µm. (F) Magnified views of magenta boxes from panel (E) acquired with STED microscopy, showing phosphorylated tau expression in dendrites. Arrows indicate phosphorylated tau expression in putative spines. Scale bar = 2 µm. See also Figure S1d & e. \*\**p* < 0.01 by two-tailed Wilcoxon test (D).

### V1 neuronal dendrites in P301S mutants are hyperactive with poor visual response tuning compared to those in wildtype controls

In order to investigate how pathological tau affects dendrites, we performed *in vivo* 2p-Ca^2+^ imaging of dendrites of layer 2/3 primary visual cortical neurons sparsely expressing GCaMP8m in P301S mutants and wildtype controls during visual stimulation (**Figure 2a; Figure S2**). Biophysical properties of thin dendrites of upper layer 2/3 cortical neurons allow axonal action potentials, encoding neuronal output, to reliably backpropagate into the entire dendritic tree^43^ (backpropagating action potentials, bAPs; **Figure 2b**) well into the distal dendrites (**Figure 2c**). Thus, calcium signals that span the dendritic tree reflect the neuronal output^43–45^. Distribution of visually responsive and non-responsive dendrites were comparable between the P301S mutants and controls (**Figure 2d**; control n = 89 measurements from 46 dendrites pooled across two imaging timepoints, 27 neurons, 6 mice, mutant dendrites n = 80 measurements from 41 dendrites pooled across two imaging timepoints, 17 neurons, 6 mice, mean ± s.e.m., visually non-responsive %: control 38.882 ± 8.224% vs. mutant 38.552 ± 10.542%, Welch’s two-tailed t test *p* = 0.9808 visually responsive %: control 61.118 ± 8.224% vs. mutant 61.448 ± 10.542%, Welch’s two-tailed t test *p* = 0.9808). The majority of visually responsive dendrites showed notable orientation tuning in both control and mutant mice with similarly shaped bAPs (**Figure 2e, f**). However, dendrites in P301S mutants were significantly more active than those in controls (**Figure 2g & h**; mean ± s.e.m., Contrast-matched normalized inferred spike counts: control n = 56 measurements from 33 dendrites pooled across two timepoints, 23 neurons, 6 mice, 1.000 ± 0.132 vs. mutant n = 53 measurements from 32 dendrites pooled across two timepoints, 14 neurons, 6 mice, 1.647 ± 0.189; two-tailed Wilcoxon test *p* = 0.0176; Contrast-matched normalized dF/F area under the curve (AUC): control dendrites 1.000 ± 0.126 vs. mutant dendrites 1.161 ± 0.083; two-tailed Wilcoxon test *p* = 0.0154). Interestingly, the visual responses of these dendrites in P301S mutants showed significantly lower tuning selectivity compared to those in controls (**Figure 2i & j**; Contrast matched normalized orientation selectivity index (OSI): mean ± s.e.m., control dendrites 1.000 ± 0.063 vs. mutant dendrites 0.781 ± 0.061; two-tailed Wilcoxon test *p* = 0.0155). The average tuning width was slightly broader for mutant dendrites. However, the difference was not statistically significant (Contrast matched normalized FWHM: mean ± s.e.m., control dendrites 1.000 ± 0.095 vs. mutant dendrites 1.069 ± 0.105; one-tailed Wilcoxon test *p* = 0.3543).

**Figure 2.**
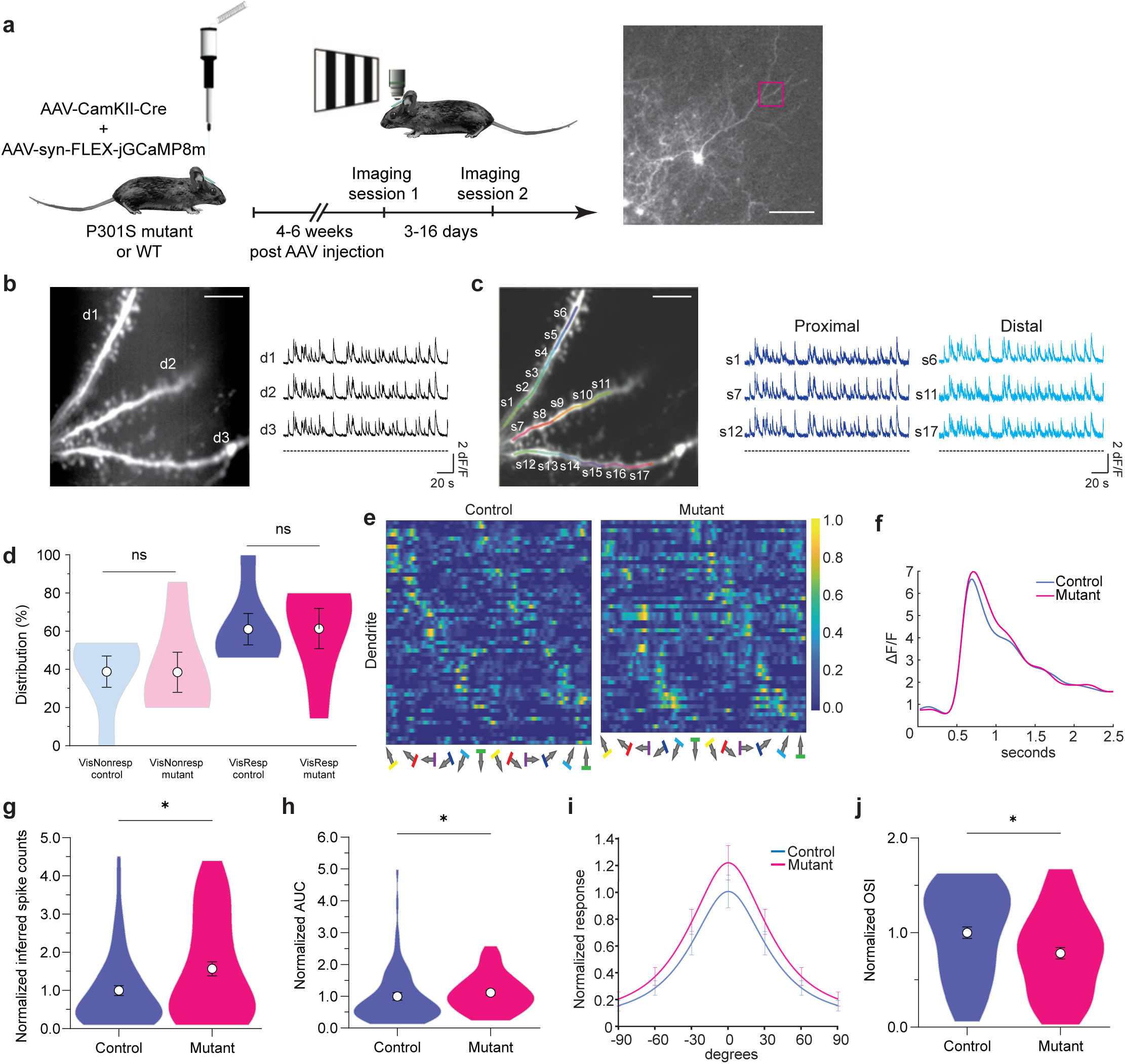
Visually-evoked dendritic responses are poorly tuned and exhibit hyperexcitability in P301S mutants. (A) Left: Schematic of the experimental design for *in vivo* two-photon calcium imaging. Ultra sparse expression of GCaMP8m expression was achieved with a combed injection of AAV-syn-FLEX-jGCaMP8m and highly dilute AAV-CamKII-Cre. Black and white drifting gratings were presented at different angles to assess orientation tuning of visual responses. Each mouse was subject to two imaging sessions. Right: An example sparsely labeled neuron expressing GCaMP8m. Scale bar = 100µm. See also Figure S2. (B) Magnified view of a dendrite boxed in panel (A), branching into three sister branches d1 - d3. The global activity shared across the three branches indicate back propagation of axonal action potentials (bAPs). Scale bar = 10µm. (C) Same dendrites as (B) but subdivided into shaft subregions s1 - s17. For each of the three sister branches, bAPs are propagated with high fidelity from proximal to distal segments. Dotted lines on the bottom of the traces indicate timing of 3s ON 1s OFF stimulus epochs. Scale bar = 10µm. (D) Distribution of visually responsive and non-responsive dendrites based on bAPs were comparable between P301S mutants and controls. (E) Heatmap of mean activity of dendrites in response to different directions of black and white moving gratings. Each response to gratings is followed by baseline activity during interleaving grey screens. Differentially oriented gratings were presented in a randomized order and sorted here for presentation. Activity values were normalized to the average peak value of the contrast-matched mutant dendrites. Only visually responsive dendrites are shown, listed by their preferred orientation in an ascending order. (F) Averaged ΔF/F bAP traces from control and mutant dendrites. (G) Total spike activity during stimulus ON periods inferred with CASCADE (ref.71 & 72) normalized to contrast-matched controls. Dendrites were hyperactive in mutants compared to controls. (control: n = 56 measurements from 33 dendrites, 23 neurons, 6 mice; mutant: n = 53 measurements from 32 dendrites, 14 neurons, 6 mice). (H) Area under the curve (AUC) of dendritic activity normalized to contrast-matched controls as a measure of activity level. Dendrites were hyperactive in mutants compared to controls. (I) Mean orientation tuning curves of all visually responsive dendrites, normalized to contrast-matched control dendrites for control and mutant mice. (J) Orientation selectivity index (OSI) normalized to contrast-matched controls. Dendrites were more poorly tuned in mutants compared to controls. \**p* < 0.05 by two-tailed Wilcoxon test.

### V1 dendritic spines in P301S mutants differ structurally and functionally from those of controls

To investigate whether the observed visually evoked hyperactivity in the P301S mutant neurons was presynaptically or postsynaptically driven, we examined the dendritic spines closely. As predicted from the hypothesized impact of tau on spine pathology^1,2,4^, we found that the spine density (number of spines per micron of dendritic shaft) was significantly lower in the dendrites of the mutants compared to those of controls (**Figure 3a**; mean ± s.e.m., control n = 84 measurements from 44 dendrites pooled across two timepoints, 27 neurons, 6 mice, 0.609 ± 0.022 vs. mutant n = 80 measurements from 41 dendrites, pooled across two timepoints, 17 neurons, 6 mice, 0.543 ± 0.014; Welch’s two-tailed t test *p* = 0.0121). When we pooled the data per neuron, the difference showed a similar trend but was no longer significant (control 0.630 ± 0.041 vs. mutant 0.576 ± 0.028; Welch’s one-tailed t test *p =* 0.1425). Thus, the observed hyperactivity in mutants was not simply caused by more abundant synaptic inputs. We next sought to examine the functional properties of the synaptic inputs arriving at the dendritic spines, using 2p-Ca^2+^ imaging-based input mapping and determined the visual response properties of the individual spines. On the dendrites we were able to successfully perform input mapping (n = 46 control dendrites from 27 neurons, 6 mice and n = 41 mutant dendrites from 17 neurons, 6 mice), we found that the distribution of visually responsive and visually non-responsive spines were comparable between mutants and controls (**Figure 3b**; Visually responsive spine percentage per dendrite; mean ± s.e.m., control n = 1694 measurements from 955 spines pooled across two timepoints, 52.090 ± 3.359 % vs. mutant n = 1422 measurements from 783 spines pooled across two timepoints 55.661 ± 4.125 %;; Visually non-responsive spine percentage per dendrite; mean ± s.e.m., control 47.911 ± 3.359 % vs. mutant 44.339 ± 4.125 %; Welch’s two-tailed t test *p* = 0.5020). When we pooled the data per neuron, the distributions remained similar and nonsignificant (Visually responsive spines per neuron; control 57.192 ± 4.001% vs mutant 50.498 ± 6.442%; Visually non-responsive spines per neuron; control 42.808 ± 4.001% vs mutant 49.502 ± 6.442%; Welch’s two-tailed t test *p* = 0.3849). As expected from rodent visual cortical neurons^46^, the preferred orientations of the synaptic inputs reflected in the dendritic spine activity were heterogeneously distributed over the length of the dendrite (**Figure 3c & d**). The majority of visually responsive spines exhibited notable orientation tuning, but not direction tuning, as indicated with the presence of peak responses at two directions separated by 180 degrees (**Figure 3e**). We proceeded to further examine visual response properties of dendritic spine activities (**Figure 3f**). When Gaussian-fit preferred orientation of all the visually responsive spines were compared to that of their visually responsive parent dendrites, distribution of relative preferred orientations was similar between the mutants and controls (**Figure 3g**; delta between spine and parental dendrite preferred orientations; mean ± s.e.m., control n = 481 measurements from 389 spines pooled across two timepoints, 32 dendrites, 22 neurons, 6 mice, 31.190 ± 1.212 degrees vs. mutant n = 587 measurements from 442 spines pooled across two timepoints, 32 dendrites, 14 neurons, 6 mice, 32.966 ± 1.117 degrees; two-tailed Wilcoxon test *p* = 0.0877) in which 37 and 35% of the spines were co-tuned with the neuronal outputs, respectively (**Figure 3h**; Kolmogorov-Smirnov two-sample test *p =* 0.2726). However, as we looked more closely, we found that the dendritic spines in the mutants showed notably poorer visual response with significantly larger tuning width compared to controls (**Figure 3i & j**; Contrast-matched normalized z-score FWHM, mean ± s.e.m: control n = 863 measurements from 675 spines pooled across two timepoints, 46 dendrites, 27 neurons, 6 mice, 1.000 ± 0.012 vs. mutant n = 790 measurements from 581 spines pooled across two timepoints, 41 dendrites, 17 neurons, 6 mice, 1.130 ± 0.023; two-tailed Wilcoxon test *p* <0.0001). The OSI was also significantly lower in mutant spines compared to controls (**Figure 3k**; Contrast-matched normalized z-score OSI, mean ± s.e.m.: control spines 1.000 ± 0.018 vs. mutant spines 0.866 ± 0.018; two-tailed Wilcoxon test *p* < 0.0001). These results mirrored our finding that the neuronal output measured by the bAP activity in the dendrites was also poorly tuned in the mutants compared to the controls (**Figure 2i & j**). Of note, the visually evoked activity of the dendritic spines in the P301S mutants were low (hypoactive) compared to those in the controls (**Figure 3l**; Contrast-matched normalized z-score AUC, mean ± s.e.m.: control spines 1.000 ± 0.020 vs. mutant spines 0.916 ± 0.015; two-tailed Wilcoxon test *p* = 0.0304). This hypoexcitability of dendritic spines observed in the mutant mice were not due to the over-subtraction of bAP signals^45^ (**Figure S3**) as the spine activity without the bAP signal removal were still significantly lower in the mutants compared to the controls (**Figure 3m**; Contrast-matched normalized z-score AUC with bAP, mean ± s.e.m.: control spines 1.000 ± 0.025 vs. mutant spines 0.899 ± 0.021; two-tailed Wilcoxon test *p* = 0.0015).

**Figure 3.**
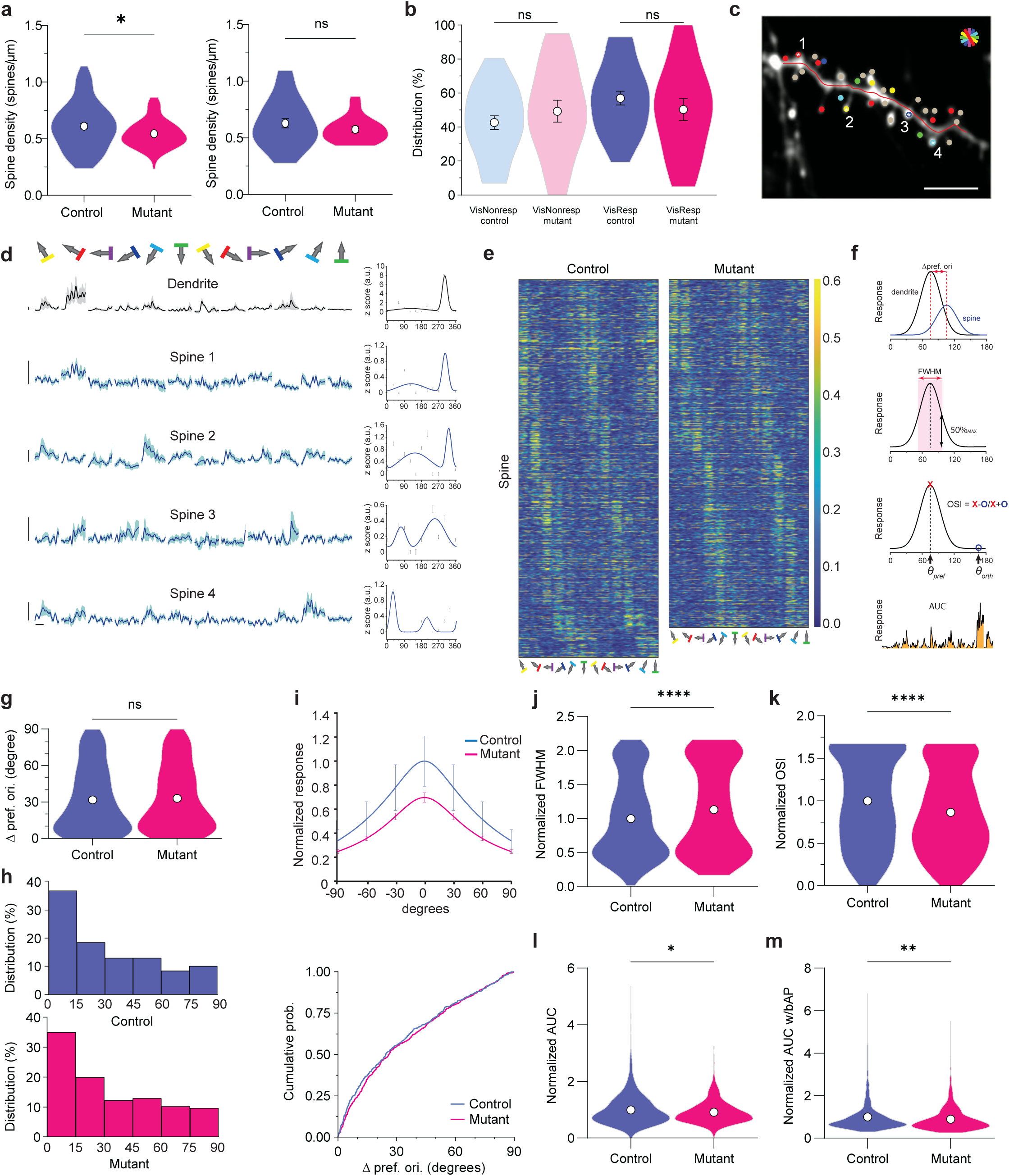
Dendritic spines and their visual response properties in P301S mice. (A) Left: Dendritic spine density per dendrite in mutants (n = 80 measurements from 41 dendrites, 17 neurons, 6 mice) compared to control mice (n = 84 measurements from 44 dendrites, 27 neurons, 6 mice). Right: Dendritic spine density per neuron in mutants (n = 17 neurons) compared to control mice (n = 27 neurons) by neuron. (B) Distribution of visually responsive and non-responsive spines were comparable between the two groups (control: n = 1694 measurements from 955 spines, 46 dendrites, 27 neurons, 6 mice; mutant: n = 1422 measurements from 783 spines, 41 dendrites, 17 neurons, 6 mice). (C) An example dendrite with their spines color coded with preferred orientation as shown in the pinwheel. Grey circles indicate visually non-responsive spines. Scale bar = 10 µm. (D) Left: Orientation tuned responses of the dendritic shaft and four representative spines. Dark line indicates the mean response across eight sweeps. Light band indicates s.e.m.. Vertical and horizontal scale bars = 2 z-scores and 1 second, respectively. Right: Direction tuning curves of the dendrite and spines 1 - 4. (E) Heatmap of mean visually responsive spine activity in response to different directions of black and white moving gratings. Each response to gratings is followed by baseline activity during interleaving grey screens. Differentially oriented gratings were presented in a randomized order and sorted here for presentation. Activity values were normalized to the average peak value of the contrast-matched control spines. Spines are organized by their preferred orientation in ascending order. The range of the color scale was capped at 0.6 to enhance visibility of the less active mutant spines. (F) Schematics of the four measurements used to characterize visual response properties of the spines: Δ preferred orientation, tuning width (FWHM), orientation selectivity (OSI), and activity level (AUC). (G) Average Δ preferred orientation compared to the parental neuronal output (dendritic bAPs) was similar between the control and mutant spines. (control: n = 481 measurements from 389 spines, 32 dendrites, 22 neurons, 6 mice; mutant: n = 587 measurements from 442 spines, 32 dendrites, 14 neurons, 6 mice). (H) Left: 37 and 35% of the spines were co-tuned with the neuronal outputs in controls and mutants, respectively. Right: Cumulative distribution of preferred orientations was comparable between the mutant and control mice. (I) Orientation tuning curves of dendritic activity normalized to contrast-matched control dendrites. (J) Tuning width (full width half max, FWHM) of the spine responses normalized to contrast-matched controls was significantly broader in mutants compared to controls. (control: n = 863 measurements from 675 spines, 46 dendrites, 27 neurons, 6 mice; mutant: n = 790 measurements from 581 spines, 41 dendrites, 17 neurons, 6 mice). (K) OSI was significantly lower in mutants compared to controls. (L) AUC normalized to contrast-matched controls was significantly lower in mutants compared to controls. (M) When measured without bAP subtraction, AUC of the spine activity remained significantly lower in mutants compared to controls. \**p* < 0.05 by two-tailed Welch’s t test (A), \**p* < 0.05, \*\**p* < 0.01, ****p < 0.0001 by two-tailed Wilcoxon test (J - M).

### Spine turnover rates were uncoupled from responsiveness in P301S mutants compared to controls

Based on our findings that 1) dendrites in P301S mutants have lower spine density (**Figure 3a**), 2) neuronal output in the mutants are hyperactive (**Figure 2g & h**), but 3) generate poorly tuned visual responses (**Figure 2j**), and 4) dendritic spine responses in mutants are hypoactive and poorly tuned (**Figures 3j-m**), we wondered whether there may be specific functional characteristics that determine the fate of the dendritic spine survival and therefore drive the observed changes in the dendritic response properties.

For the spines whose identities were successfully tracked across two timepoints, the fate of the dendritic spine survival was recorded after the second timepoint, denoting the spines as lost, gained, or retained, respectively (**Figure 4a**). The initial assessment of the spine turnover performed using our custom analysis software, AUTOTUNE^45^, was further evaluated by trained experts who were blind to the genotype of the subjects. The two assessments were in agreement for the most part, however, corrections by experts were sometimes needed, especially for cases in which there were multiple spines in close proximity or the position of the spine head has moved slightly (**Figure S4**).

**Figure 4.**
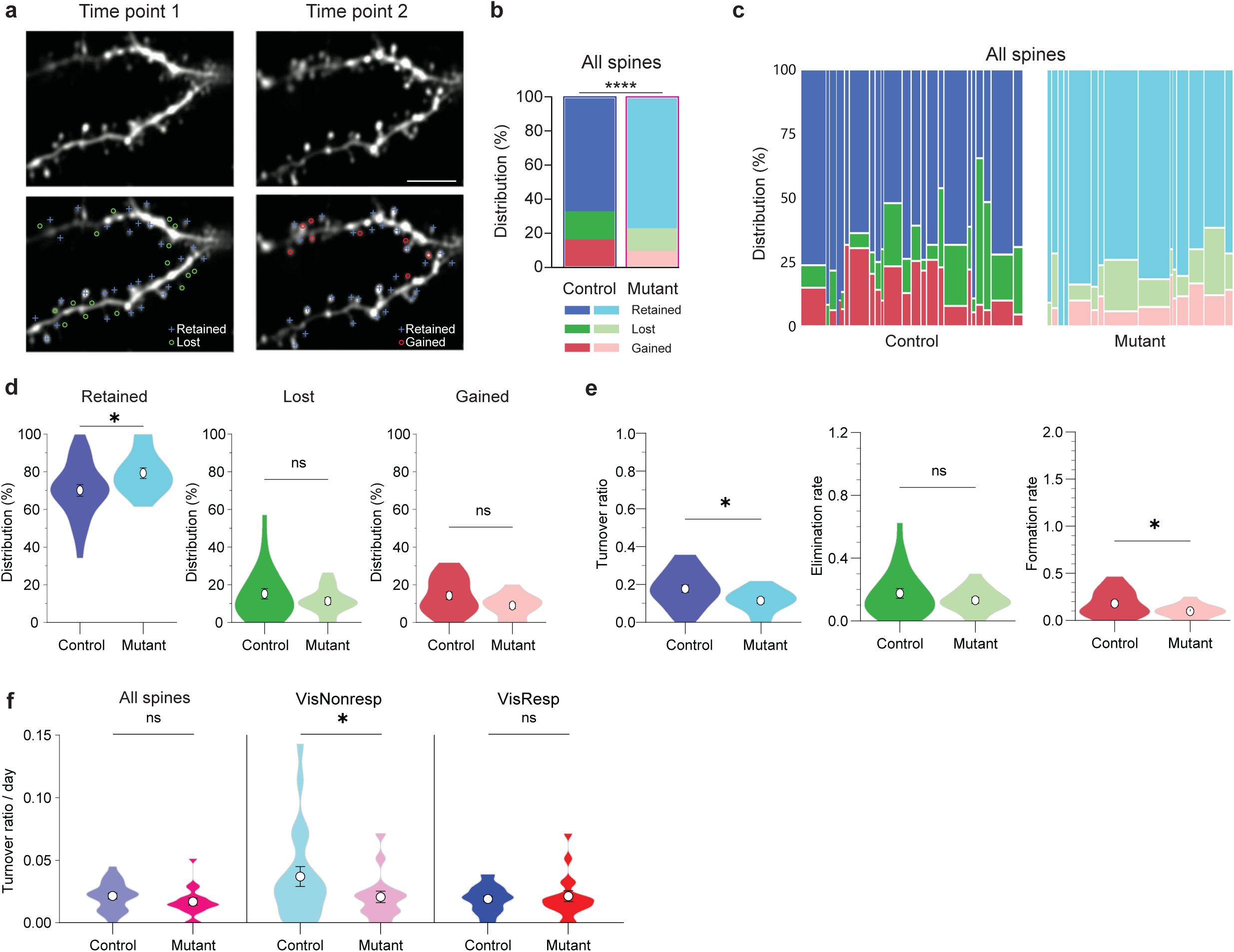
Dendritic spine turnover in P301S mice. (A) Example dendrites imaged at two timepoints separated by 7 days. Blue cross, green circle, and red circle indicate those spines that were retained, lost, and gained across the two imaging sessions, respectively. Scale bar = 10µm. See also Figure S4. (B) Distribution of spine turnover in all imaged spines. Blue/light blue, green/light green, and red/pink bars indicate retained, lost, and gained spines in control and mutants, respectively. (control: n = 646 retained, 161 lost, 153 gained spines from 43 dendrites, 24 neurons, 6 mice; mutant: n = 626 retained, 109 lost, 74 gained spines from measurements from 41 dendrites, 17 neurons, 6 mice). (C) Distribution of spine turnover by neuron. Each column represents a neuron ordered from left to right by the length of interval between imaging sessions for each genotype, and the relative width reflects proportionate spine counts per neuron. (D) Comparison of percentage of retained (left; dark blue vs. light blue), lost (middle; dark green vs. light green), and gained spines (right; red vs. pink) among all spines per neuron between controls and mutants. See also Figure S4. (E) Comparison of spine turnover ratio (left; dark blue vs. dark pink), elimination rate (middle; dark green vs. light green), and formation rate (right; red vs. pink) among all spines per neuron between controls and mutants. (F) Comparison of spine turnover ratio per neuron per day which accounts for the difference in the length of interval between the two imaging timepoints shown for all spines (left), visually non-responsive spines only (middle), and visually responsive spines only (right). (Visually non-responsive spines, control: n = 320 retained, 83 lost, 82 gained spines from 40 dendrites, 22 neurons, 6 mice; mutant: n = 288 retained, 47 lost, 28 gained spines from measurements from 33 dendrites, 16 neurons, 6 mice; Visually responsive spines, control: n = 326 retained, 78 lost, 70 gained spines from 43 dendrites, 24 neurons, 6 mice; mutant: n = 338 retained, 62 lost, 46 gained spines from measurements from 39 dendrites, 17 neurons, 6 mice). \*\*\*\**p* < 0.0001 by Chi-Square test (B). \**p* < 0.05 by two-tailed Wilcoxon test (D), two-tailed Welch’s t-test (E), and one-tailed Wilcoxon test (F).

We found that a significantly larger proportion of the spines were being retained across the two timepoints in mutant mice. This spine turnover pattern was evident in two different distribution analyses that compared between the genotypes. The first analysis calculated the % distribution of spine turnover status across all imaged spines (**Figure 4b**; % retained, lost or gained spines; control spines n = 960 total from 43 dendrites, 24 neurons, 6 mice; 67.292, 16.771, 15.938% vs. mutant spines n = 809 total from 41 dendrites, 17 neurons, 6 mice; 77.380, 13.473, 9.147%; Chi-Square test *p* < 0.0001). The second analysis assessed the distribution per neuron (**Figure 4c & d**; % retained per neuron control n = 24 neurons, 646 spines, 70.375 ± 3.098 vs mutant n = 17 neurons, 626 spines, 79.500 ± 2.837, two-tailed Wilcoxon *p* = 0.0415; % lost per neuron control n = 161 spines, 15.272 ± 2.806% vs. mutant n = 109 spines, 11.471 ± 1.850%, two-tailed Wilcoxon *p* = 0.7590; % gained per neuron control n = 153 spines, 14.353 ± 1.925 vs. mutant n = 74 spines, 9.029 ± 1.549, two-tailed Wilcoxon *p* = 0.1360). Interestingly, this difference in the spine turnover proportion seems to be driven by a larger proportion of visually non-responsive spines that were retained in the mutants (**Figure S4c-e**; Visually non-responsive spines % retained per neuron, control n = 320 spines from 42 dendrites, 23 neurons, 6 mice, 58.402 ± 6.500% vs. mutant n = 288 spines from 33 dendrites, 16 neurons, 6 mice, 76.911 ± 5.021%, two-tailed Wilcoxon *p* = 0.0366; % lost per neuron control n = 83 spines, 18.441 ± 5.233% vs. mutant n = 47 spines, 13.446 ± 3.685%, two-tailed Wilcoxon *p* = 0.9438; % gained per neuron control n = 83 spines, 23.157 ± 5.467% vs. mutant n = 28 spines, 9.643 ± 2.512%, two-tailed Wilcoxon *p* = 0.2483; Visually responsive spines % retained per neuron, control n = 326 spines from 43 dendrites, 24 neurons 73.411 ± 2.827% vs. mutant n = 338 spines from 39 dendrites, 17 neurons, 75.395 ± 3.618%, two-tailed Wilcoxon *p* = 0.8360; % lost per neuron control n = 78 spines, 12.952 ± 2.911% vs. mutant n = 62 spines, 14.381 ± 3.307%, two-tailed Wilcoxon *p* = 0.4137; % gained per neuron control n = 70 spines, 13.638 ± 1.895% vs. mutant n = 46 spines, 10.224 ± 2.584%, two-tailed Wilcoxon *p* = 0.2025).

To supplement this analysis, we have additionally computed spine formation rate, elimination rate, as well as the turnover ratio (see Methods). The spine turnover ratio and the spine formation rate were significantly larger in controls compared to the mutants whereas the spine elimination rate was similar between the two groups (**Figure 4e**; control n = 24 neurons, mutant n = 17 neurons, spine turnover ratio, control 0.179 ± 0.020 vs. mutant 0.116 ± 0.015, Welch’s two-tailed t test *p* = 0.0139; spine formation rate, control 0.182 ± 0.028 vs. mutant 0.100 ± 0.017, Welch’s two-tailed t test *p* = 0.0162; spine elimination rate, control 0.177 ± 0.031 vs. mutant 0.132 ± 0.019, two-tailed Wilcoxon test *p* = 0.7009). While the average interval between the two timepoints was not statistically different between the genotypes, (control 8.79 ± 0.69 vs. mutant 7.93 ± 1.24, two-tailed Wilcoxon p = 0.4689), the interval ranged from 3 to 16 days. Thus, we further examined the turnover ratio per day. The spine turnover ratio per day showed the similar pattern in which the ratio is larger in controls compared to the mutants. Interestingly, this observation was more prominent among those spines that were visually non-responsive (**Figure 4f**; all spines included, spine turnover ratio per neuron per day, control n = 24 neurons, 0.021 ± 0.002 vs. mutant n = 17 neurons, 0.017 ± 0.003, one-tailed Wilcoxon *p* = 0.0657; visually non-responsive spines, control = 22 neurons, 0.037 ± 0.008 vs. mutant n = 16 neurons, 0.021 ± 0.005, one-tailed Wilcoxon *p* = 0.0433; visually responsive spines, control n = 24 neurons, 0.019 ± 0.002 vs. mutant n = 17 neurons, 0.021 ± 0.004, one-tailed Wilcoxon *p* = 0.3295).

### Structure-function relationships in the maintenance of dendritic spines is disrupted in P301S mutants

Probing for functional underpinnings of the poor visual response tuning selectivity we observed in the mutant dendrites, we examined whether activity profiles of the spines directed their fate in spine turnover. Surprisingly, in both control and mutant mice, stably retained spines were least active of all spines (**Figure 5a**; AUC normalized to those of retained spines per dendrite; control mice, mean ± s.e.m.: retained spines n = 646, 1.000 ± 0.020 lost spines n = 161, 1.104 ± 0.059, gained spines n = 153, 1.080 ± 0.049 from 46 dendrites, 27 neurons, 6 mice; Dunn’s multiple comparison test, retained vs. lost, retained vs. gained, lost vs. gained, *p* = 0.0200, *p* > 0.9999, *p* = 0.5300, respectively; mutant mice, mean ± s.e.m.: retained spines n = 626, 1.000 ± 0.016, lost spines n = 109, 1.165 ± 0.041, gained spines n = 74, 1.019 ± 0.050 from 41 dendrites, 17 neurons, 6 mice; Dunn’s multiple comparison test, retained vs. lost, retained vs. gained, lost vs. gained, *p* < 0.0001, *p* > 0.9999, *p* = 0.0303, respectively). Despite this observation, the retained spines were significantly larger in size than lost or gained spines in control dendrites whereas spine sizes were more comparable regardless of the turnover fate in mutants (**Figure 5b**; Spine size normalized to those of retained spines per dendrite; control mice, mean ± s.e.m.: retained spines 1.000 ± 0.0170, lost spines 0.849 ± 0.033, gained spines 0.899 ± 0.037; Dunn’s multiple comparison test, retained vs. lost, retained vs. gained, lost vs. gained, *p* = 0.0005, *p* = 0.0406, *p* > 0.9999, respectively; mutant mice, mean ± s.e.m.: retained spines 1.000 ± 0.016, lost spines 0.968 ± 0.038, gained spines 0.954 ± 0.047; Dunn’s multiple comparison test, retained vs. lost, retained vs. gained, lost vs. gained, *p* = 0.8939, *p* > 0.9999, *p* > 0.9999, respectively).

**Figure 5.**
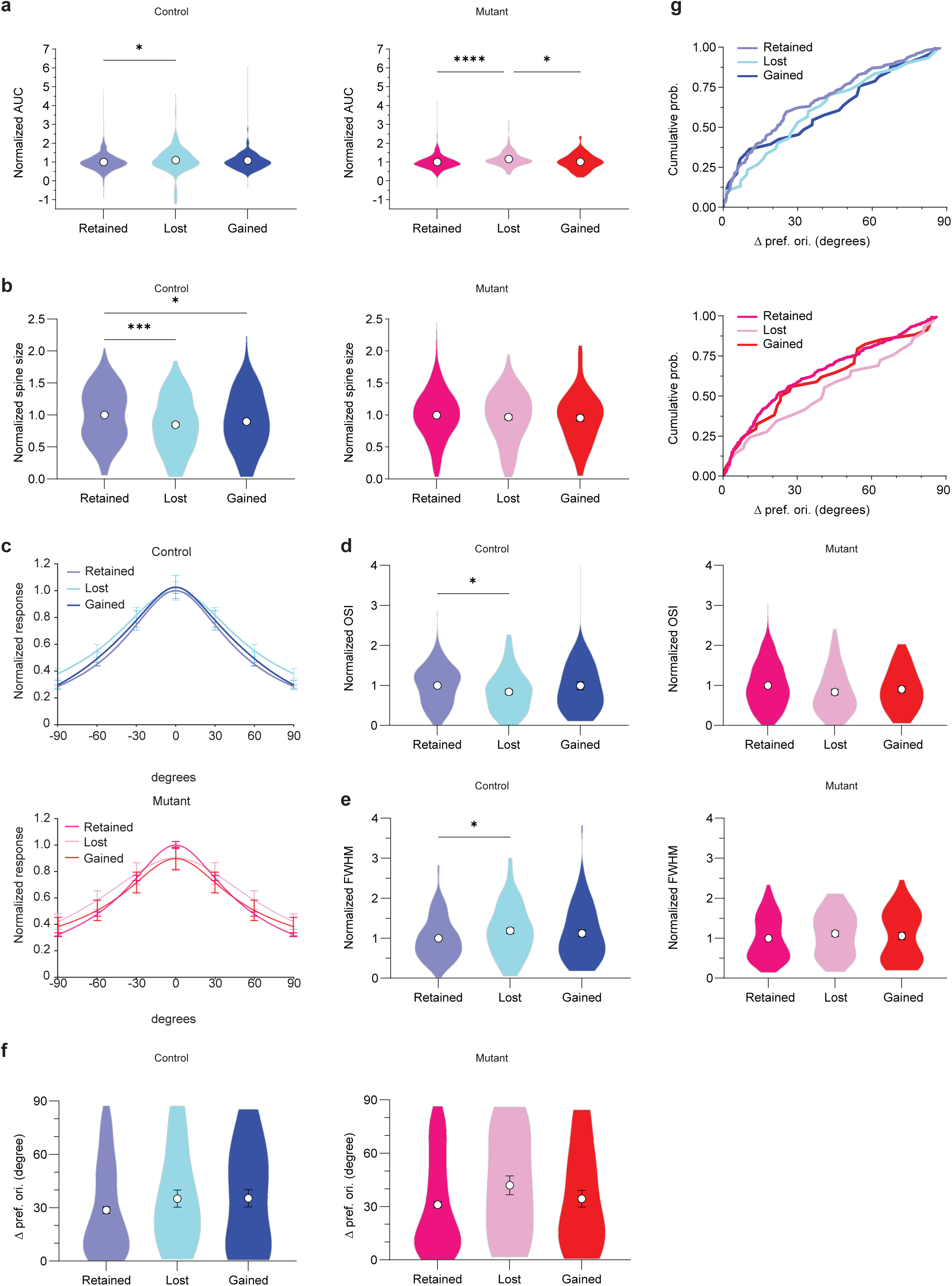
Visual response properties of the spines and their fate in turnover. (A) AUC normalized to retained spines per dendrite shown for retained, lost, and gained spines from control and mutant mice. (control: n = 646 retained, 161 lost, 153 gained spines from 43 dendrites, 24 neurons, 6 mice; mutant: n = 626 retained, 109 lost, 74 gained spines from measurements from 41 dendrites, 17 neurons, 6 mice). (B) Spine size normalized to retained spines per dendrite shown for retained, lost, and gained spines from control and mutant mice. (C) Orientation tuning curves normalized to retained spines per dendrite shown for retained, lost, and gained spines from control and mutant mice. (D) OSI normalized to retained spines per dendrite shown for retained, lost, and gained spines from control and mutant mice. (control: n = 326 retained, 78 lost, 70 gained spines from 43 dendrites, 24 neurons, 6 mice; mutant: n = 338 retained, 62 lost, 46 gained spines from measurements from 39 dendrites, 17 neurons, 6 mice). (E) FWHM normalized to retained spines per dendrite shown for retained, lost, and gained spines from control and mutant mice. (F) Difference in the individual spine’s preferred orientation and that of the neuronal output (bAP tuning of the parent dendrite; Ä preferred orientation) shown for retained, lost, and gained spines from control and mutant mice. (control: n = 168 retained, 30 lost, 33 gained spines from 30 dendrites, 20 neurons, 6 mice; mutant: n = 256 retained, 29 lost, 34 gained spines from measurements from 32 dendrites, 14 neurons, 6 mice). (G) Cumulative probability functions showing Δ preferred orientation shown for retained, lost, and gained spines from control and mutant mice. See also Figure S5. \**p* < 0.05, \*\*\**p* < 0.001, ****p < 0.0001 by two-tailed Wilcoxon test (A, B, D, E).

When we compared the stimulus response tuning across all visually responsive spines (**Figure 5c**), we found that in control mice, retained spines were more orientation selective with significantly higher OSI compared to lost spines, a trend that was there but did not reach a statistical significance in the mutants (**Figure 5d**; Contrast-matched z-score OSI normalized to retained spines per dendrite, mean ± s.e.m.: control mice, retained spines n = 326, 1.000 ± 0.027, lost spines n = 78, 0.842 ± 0.058, gained spines n = 70, 0.994 ± 0.099 from 43 dendrites, 24 neurons, 6 mice; Dunn’s multiple comparison test, retained vs. lost, retained vs. gained, lost vs. gained, *p* = 0.018, *p* = 0.709, *p* = 0.740, respectively; mutant mice, retained spines n = 338, 1.000 ± 0.031, lost spines n = 62, 0.834 ± 0.071, gained spines n = 46, 0.902 ± 0.075 from 39 dendrites, 17 neurons, 6 mice; Dunn’s multiple comparison test, retained vs. lost, retained vs. gained, lost vs. gained, *p* = 0.0648, *p* = 0.9925, *p* > 0.9999, respectively). In addition, retained spines in controls exhibited significantly sharper tuning compared to lost spines (**Figure 5e**; Contrast-matched z-score FWHM normalized to retained spines per dendrite, mean ± s.e.m: control mice, retained spines 1.000 ± 0.030, lost spines 1.149 ± 0.074, gained spines 1.162 ± 0.091; Dunn’s multiple comparison test, retained vs. lost, retained vs. gained, lost vs. gained, *p* = 0.0482, *p* > 0.9999, *p* = 0.8378, respectively. This pattern was not observed in mutant mice (mean ± s.e.m: mutant mice, retained spines 1.000 ± 0.030, lost spines 1.117 ± 0.062, gained spines 1.057 ± 0.092; Dunn’s multiple comparison test, retained vs. lost, retained vs. gained, lost vs. gained, *p* = 0.3869, *p* > 0.9999, *p* > 0.9999, respectively).

No obvious pattern was observed in the preferred orientations exhibited by the retained, lost, or gained spines relative to that of the parental dendrite bAPs in the mutants or the controls (**Figure 5f**; delta preferred orientation angle compared to parental dendritic tuning, mean ± s.e.m.: visually tuned spines that were found on visually tuned dendrites, control mice, retained spines n = 168, 28.643 ± 1.926 degrees, lost spines n = 30, 35.147 ± 4.808, gained spines n = 33, 35.344 ± 4.933 from 30 dendrites, 20 neurons, 6 mice; Dunn’s multiple comparison test, retained vs. lost, retained vs. gained, lost vs. gained, *p* = 0.5389, *p* = 0.8387, *p* >0.9999, respectively; mutant mice, retained spines n = 256, 31.041 ± 1.631, lost spines n = 29, 41.985 ± 5.327, gained spines n = 34, 34.443 ± 4.715 from 32 dendrites, 14 neurons, 6 mice; Dunn’s multiple comparison test, retained vs. lost, retained vs. gained, lost vs. gained, *p* = 0.1625 *p* > 0.9999, *p* = 0.9699, respectively). The cumulative distribution function of the preferred orientation of the spines did not differ among the turnover status in either controls or mutants. (**Figure 5g**; Kolmogorov-Smirnov test, retained vs. lost, retained vs. gained. lost vs. gained in controls *p =* 0.2542*, p =* 0.2461*, p =* 0.7365; in mutants *p =* 0.1392*, p =* 0.7829*, p =* 0.4705). The average preferred orientations compared to the neuronal output for retained, lost, or gained spines were not statistically different between control and mutant mice (**Figure S5**; delta preferred orientation angle compared to parental dendritic tuning, control vs. mutant, Dunn’s multiple comparison test, retained spines *p* = 0.9629; lost spines *p* > 0.9999; gained spines *p* > 0.9999).

### Trained classifier models identify dendritic activity profile over spine density as key predictor that discriminates P301S mutants from controls

Using five dendrometric predictors described above, global bAP-inferred spiking activity in the dendrites, AUC, OSI, FWHM, and spine density, we asked if these features were sufficient for training classifier models to reliably discriminate between the genotypes. Of the 34 binary classification learners we trained (see Methods), we selected three models that reliably and consistently discriminated mutants from controls across eight repeated trials: Gaussian Naïve Bayesian, Quadratic Discriminant, and Quadratic Support Vector Machine (SVM). Receiver Operating Characteristic (ROC) analyses of these trained models produced curves with AUC of 0.8091, 0.8364, and 0.8273 with test accuracy of 76.2, 76.2 and 76.2%, exhibiting good discriminability (**Figure 6a**). In order to determine the influence of each predictor, we calculated permutation feature importance (PFI) across 10 permutations. Of the five predictors, metrics of dendritic activity such as inferred spiking activity and AUC exhibited consistently high PFI values followed by response tuning metrics, FWHM and OSI. By contrast, spine density which is a structural metric ranked with the lowest PFI in all three models, indicating that functional rather than structural parameters may better define the impact of tauopathy at this stage during an early onset of pathology (**Figure 6b**; PFI for AUC, inferred total spikes, FWHM, OSI, and spine density, mean ± s.e.m., Quadratic Discriminant: 0.167 ± 0.016, 0.148 ± 0.023, 0.0714 ± 0.022, 0.000 ± 0.021,-0.024 ± 0.016, respectively. Statistical significance by Dunn’s multiple comparison tests were found between AUC vs. spine density, inferred total spikes vs. spine density, AUC vs. OSI, and inferred total spikes vs. OSI; *p* = 0.0001, 0.0009, 0.0012, and 0.0078; Quadratic SVM: 0.143 ± 0.012, 0.062 ± 0.019, 0.076 ± 0.020, 0.038 ± 0.032, 0.014 ± 0.010, respectively.

**Figure 6.**
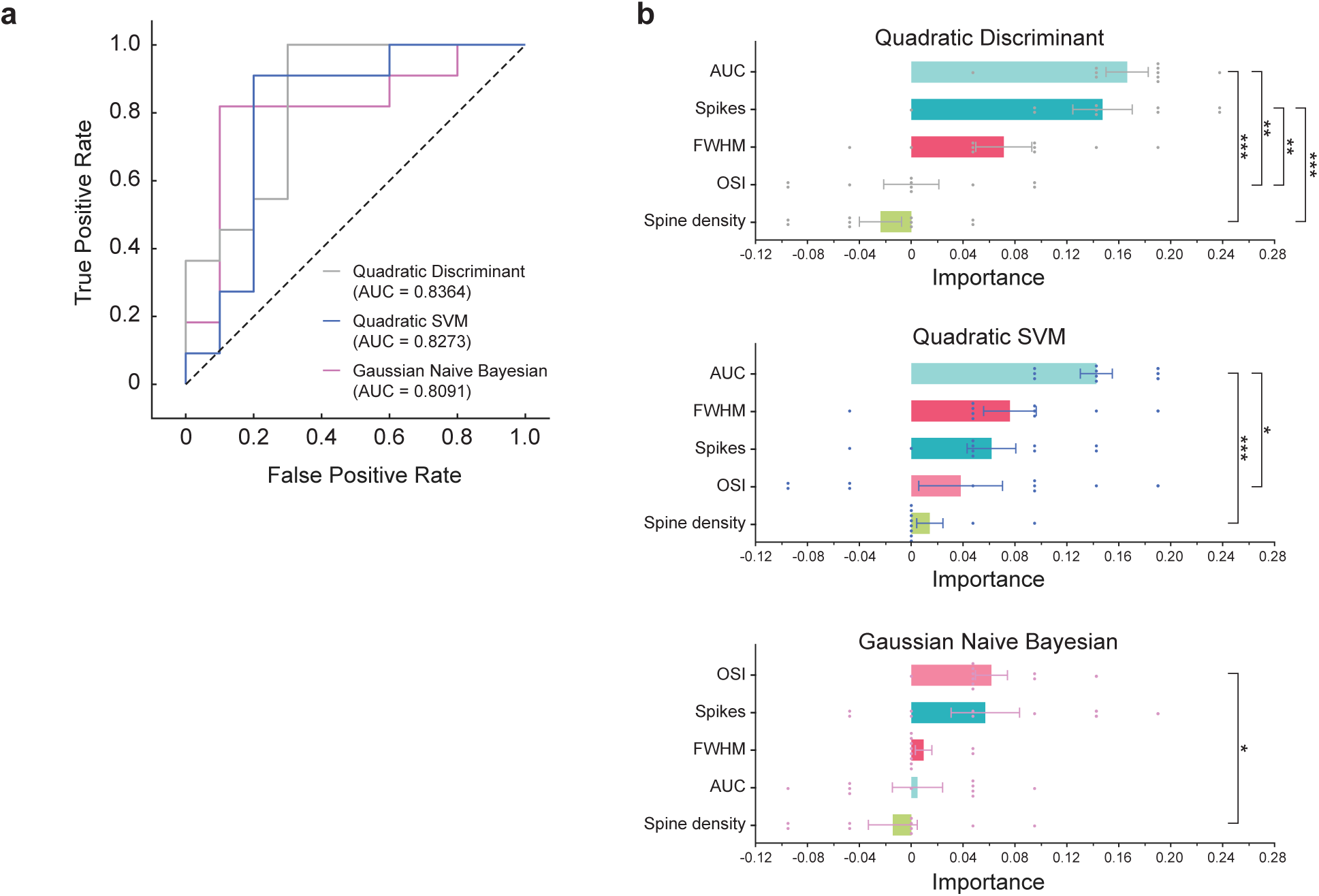
ROC analyses of classifier models trained with five dendrometric parameters, AUC, inferred spike counts, spine density, FWHM, and OSI to test discriminability between mutants and controls. (B) Three select models, Quadratic Discriminant, Quadratic SVM, and Gaussian Naïve Bayesian were trained and tested. All three models successfully discriminated between the genotypes. (C) Permutation importance of each of the parameters shown for the three models.

Statistical significance by Dunn’s multiple comparison tests were found between AUC vs. spine density and AUC vs. OSI; *p* = 0.0004 and 0.0336, respectively; Gaussian Naïve Bayesian: 0.005 ± 0.019, 0.057 ± 0.026, 0.010 ± 0.006, 0.062 ± 0.012,-0.014 ± 0.02, respectively. Statistical significance by Dunn’s multiple comparison tests were found between OSI vs. spine density; *p* = 0.0281).

## DISCUSSION

Using *in vivo* longitudinal 2p-Ca^2+^ imaging of dendritic shaft and spine activity with GCaMP8m, we found that neuronal output tuning of layer 2/3 visual cortical neurons in P301S control mice was well-tuned but less active, indicating a high precision in their visual response. This is in contrast to the neurons in P301S mutants whose outputs were poorly tuned and aberrantly hyperactive (**Figure 2e-g**). Intriguingly, this hyperactivity of the neurons in the mutants did not result from excessive number of synaptic inputs. In fact, the spine density was reduced in the mutant dendrites (**Figure 3a**). This combination of hyperactivity and decreased spine density is reminiscent of dendritic structural degeneration that was observed in the hyperactive hippocampal CA1 pyramidal neurons in a mouse model of AD, APP/PS1^47^. Moreover, the dendritic spine activity in response to visual inputs was rather reduced in the mutants. Thus, visually responsive neuronal outputs in the tauopathy mice were hyperactive despite the reduced dendritic spine response.

A few possible scenarios may provide explanation to these paradoxical observations: 1) Pathological tau in the dendritic compartments could dampen the postsynaptic response to glutamate. Indeed, the increased concentration of tau in the dendritic spines is associated with the removal of AMPA and NMDA receptors from the postsynaptic density^21^. The exclusion of NMDA receptors from the synapses may also explain the deficits in plasticity observed in the visual cortex of another mouse model of tauopathy reported by others^48^. 2) The inefficient dendritic spine response could also be due to structural weakening of the synapses. A mismatched structural instability of the presynaptic terminals and postsynaptic spines during the early stages of tauopathy has been reported^49^. These structural changes could be driven by either axonal or dendritic aggregation of pathological tau or both. Since we could already observe the dendritic spine loss in the mutant visual cortex, the remaining spines could also be undergoing mismatched structural changes at the synapse, resulting in reduced dendritic spine response. 3) The reduced spine activity in combination with poor tuning could result from abnormal glutamate spillover. Pathological tau has been linked to a 40% reduction in the expression of glutamate transporters^50^. Insufficient reuptake of glutamate may cause nonspecific activation^51,52^ and compromise the input specificity of the neuronal output. The resultant aberrant synaptic activation may trigger compensatory homeostatic downscaling of synaptic strength^53^ (circling back to scenario 1). Finally, 4) there could be a pathological mechanism at play in postsynaptic dendritic nonlinear integration that abnormally amplifies synaptic inputs. Ca^2+^ conductance plays an essential role in dendritic integration in many ways. It supports dendritic NMDA receptor spikes^54^ upon arrival of strong inputs that trigger a cascade of voltage-dependent events, including firing of dendritic spikes^43^ and internal store Ca^2+^ release from endoplasmic reticulum^55^ (ER). ER and mitochondria are key organelles that regulate Ca^2+^ dynamics in the neuronal dendrites. Their functions directly affect synaptic integration and neuronal output^56^, and their size and distribution along the dendrites would impact the spread, spatial pattern, and input-specificity of local dendritic response. Of note, an increased Ca^2+^ release from ER as well as mitochondrial dysfunction have been reported as early markers of neurodegenerative diseases that precedes the onset of cognitive decline^57,58^, and mounting evidence have highlighted the tight coupling between mitochondrial dysfunction in tauopathies such as abnormal mitochondrial morphology^59–61^. Whether ER or mitochondrial deficits, and their impact on Ca^2+^ regulation, directly lead to an increase in dendritic excitability remains to be tested.

Consistent with the poor visual response tuning of neuronal outputs found in mutant mice, response tuning of individual spines was also poor compared to those in controls (**Figure 3j-m**). Because overall decreased quality in visual tuning on both presynaptic inputs and postsynaptic outputs would only provide snapshots of dendritic spine functions in mutants, we decided to track potential changes in dendritic spine functions across days through spine turnover. We found that spines receiving well-tuned synaptic inputs were preferentially stabilized while those receiving noisy inputs were turned over in control mice, with retained spines exhibiting greater tuning selectivity than the spines that were lost before the second timepoint (**Figure 5a, d, e**). Moreover, in control mice, retained spines were significantly larger in size compared to those that were lost or gained. These findings illustrate a scenario in which maintenance of well-tuned spines contribute to the overall preservation of tuned neuronal output. This pattern was notably absent from the visual neurons of the tauopathy mice in which dendritic spines on average were sparser and poorly tuned with reduced activity levels (**Figure 3a, j-m**), and the spine size did not differ among retained, lost, and gained spines (**Figure 5b**).

The three classifier models we selected were able to successfully discriminate between the genotypes of the dendrites based on just five simple metrics, spiking activity, AUC, OSI, FWHM, and spine density. Of the metrics we examined, neurofunctional metrics such as spiking activity, AUC and tuning were better predictors than spine density. Our findings from the classifier models may indicate that a shift in visual response functions precedes the structural degeneration of the spines at least in the superficial layer 2/3 of the cortex, rather than the structural loss of the spines driving the functional deterioration. Taken together, these findings demonstrate functional impact of tauopathy on cortical visual processing at both presynaptic input and postsynaptic integration levels. Interestingly, layer 4 neurons exhibited reduced deposition of hyperphosphorylated tau compared to layer 2/3 or layer 5 neurons (**Figure S1b & c**). Whether this input layer is functionally spared or how the tau pathology progresses through the cortical layers remains unclear.

Our longitudinal study paints a model in which cortical visual processing is slowly affected in progressive tauopathy. Indeed, visual impairments are a common symptom observed in many neurodegenerative diseases with concomitant tauopathy^26–32^. As such, early onset of visual impairments has garnered attention as an indicator of preclinical neurodegenerative diseases^33–35^. Importantly, a six-week program of visual rehabilitation was reported effective in improving not only visual functions, but also cognitive measures including logical memory in elderly participants with mild cognitive deficits^62^. Repeated presentation of grating stimulus set to a single orientation is known to exert experience-dependent response enhancement to that stimulus in the mouse visual cortex^63^. Based on our finding that spines that survive the spine turnover show better tuning, response enhancement through simple visual rehabilitation may exert spine protective effects. It remains to be seen whether gained spines may also be better retained by visual training.

### Limitations of the study

In this study, we found a decrease in the dendritic spine density in P301S mutant mice. While this is in line with the neurodegenerative nature of the mutation^64^, it is possible that our use of GCaMP8m in vivo dynamic fluorescence, rather than a static fluorescence like those from eGFP in fixed tissues, could have overlooked small, less active spines. However, because we produce spine region of interest (ROI) map on the raw image stack before bAP subtraction, the bAP will invade majority of the spines, making them detectable despite potential difference in the level of their individual spine-specific activities. Moreover, the spine density and the daily turnover ratio observed in our control mice are in line with the previous report that employed a static fluorescence of eGFP expressed in mouse V1 neurons^65^ although the latter was slightly lower in our study at 2.1% which may be explained by the old age span used in the study. Nonetheless, future studies employing spine density and turnover analyses using a static fluorescence in fixed tissues would provide a definitive answer.

## RESOURCE AVAILABILITY

### Lead contact

Requests for further information and resources should be directed to and will be fulfilled by the lead contact, Ikuko Smith.

### Materials availability

This study did not generate new unique reagents. Materials used in this study are commercially available.

### Data and code availability

- All data reported in this paper will be shared by the lead contact upon request. Processed data of calcium imaging have been deposited at Mendeley Data and are publicly available as of the date of publication. The DOI is listed in the key resources table.
- All original code has been deposited at GitHub and is publicly available at https://github.com/gtzook/iSLAB-data-analysis as of the date of publication.
- Any additional information required to reanalyze the data reported in this paper is available from the lead contact upon request.

## ACKNOWLEDGMENTS

This work was supported by Brain Research Foundation Fay Frank Seed Grant (BRFSG-2019-05), Brain and Behavior Research Foundation NARSAD Young Investigator Grant (30481) and the NIH (NINDS R01NS128079) to I.T.S.. We thank Dr. Ramon Velazquez at Arizona State University for kindly providing us with PS19 line founder breeders. We thank the members of the I. Smith lab and Dr. Spencer Smith for providing valuable comments on the earlier versions of this manuscript and Gabriel Zook for his illustrations. We also thank Andy Alexander and the NRI microscopy core director, Ben Lopez, for sharing their equipment and expertise.

## AUTHOR CONTRIBUTIONS

Conceptualization, L.M.A. and I.T.S.; methodology, L.M.A, K.C., and I.T.S.; Investigation, L.M.A. and K.C.; writing—original draft, L.M.A., K.C., and I.T.S.; writing—review & editing, L.M.A., K.C., and I.T.S.; funding acquisition, I.T.S.; resources, I.T.S.; supervision, I.T.S.

## DECLARATION OF GENERATIVE AI AND AI-ASSISTED TECHNOLOGIES

No AI or AI-assisted technologies were used during the preparation of this manuscript.

## SUPPLEMENTAL INFORMATION

**Document S1. Figures S1–S6**

## STAR★METHODS

### KEY RESOURCES TABLE

Key resources table

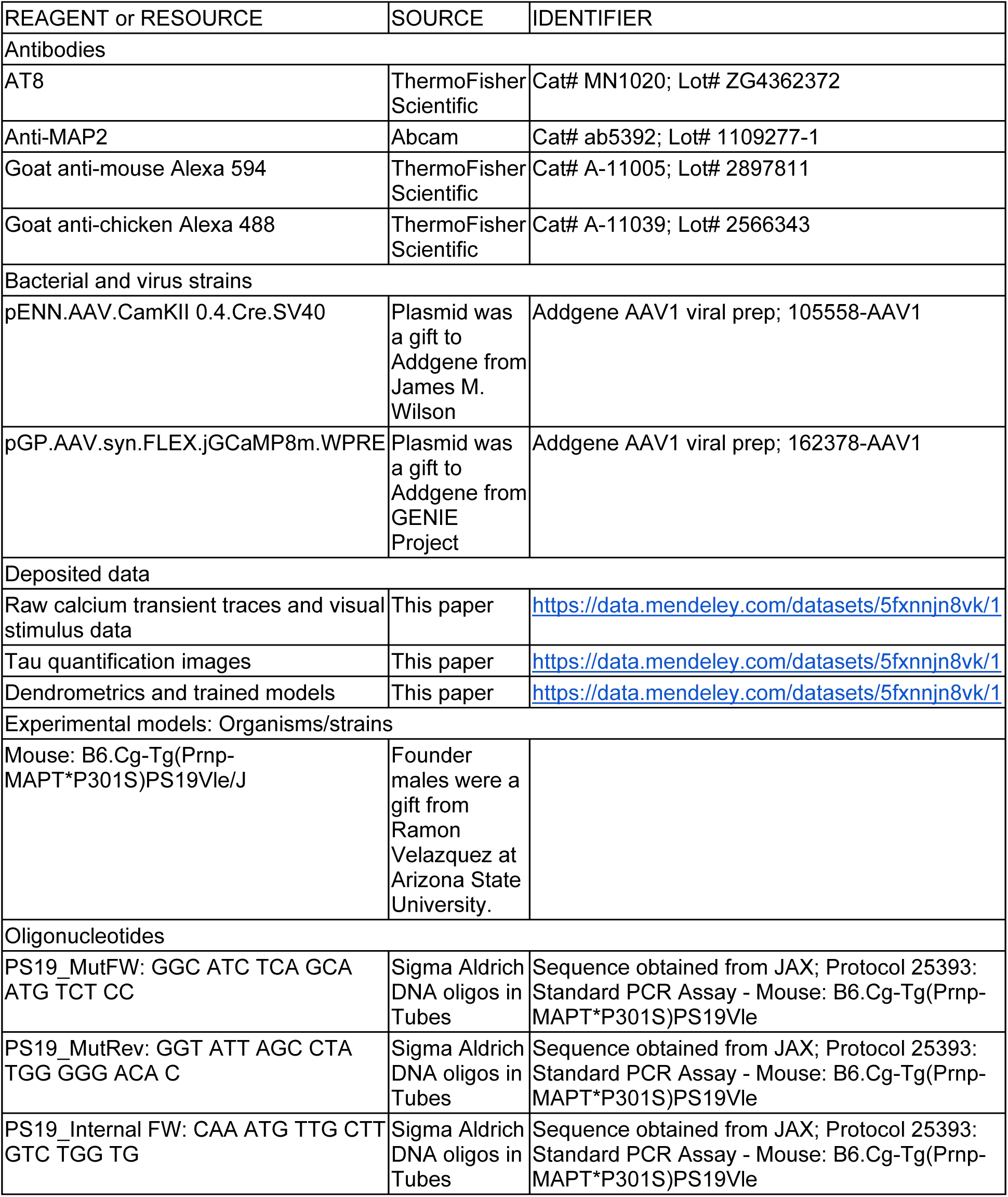
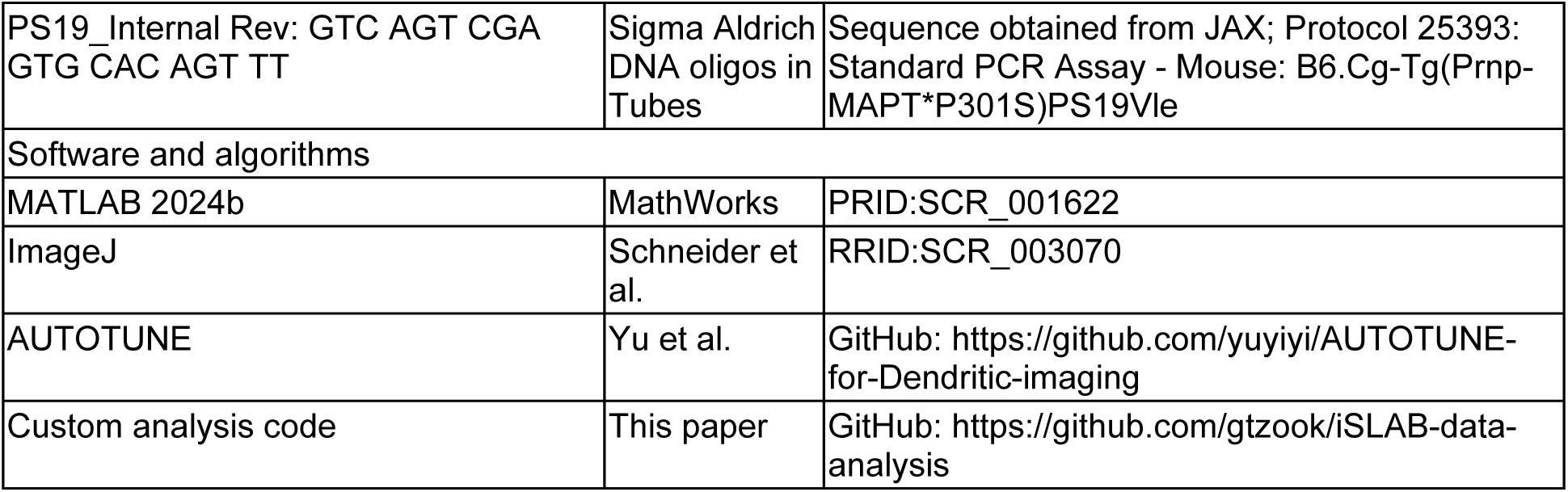

### EXPERIMENTAL MODEL AND STUDY PARTICIPANT DETAILS

All procedures involving mice were performed in accordance with the guidelines and regulations of the US Department of Health and Human Services and approved by the Institutional Animal Care and Use Committee at University of California, Santa Barbara. Adult P301S mice (PS19 line) and non-carrier wildtype controls of both sexes between 5 months and 10 months of age (founder breeders kindly provided by Dr. Ramon Velazquez at Arizona State University) were housed under a 12h/12h dark-light reversed cycle with *ad libitum* access to food and water. P301S transgenic mice express mutant tau consisting of four microtubule-binding domains and one N-terminal insert (4R/1N) that harbors disease-associated P301S missense mutation, whose expression is driven by the mouse prion protein promoter and renders the tau protein susceptible to hyperphosphorylation. This transgene causes a 249 Kb deletion at the insertion site on chromosome 3 which spares any known genes^66^. Mice were ear clipped around 24 days of age and genotyped by PCR. Control and mutant were non-littermates from the same colony. Age-matched mice were used for functional and immunohistochemical analyses (Table 1 & 2).

**Table 1.**
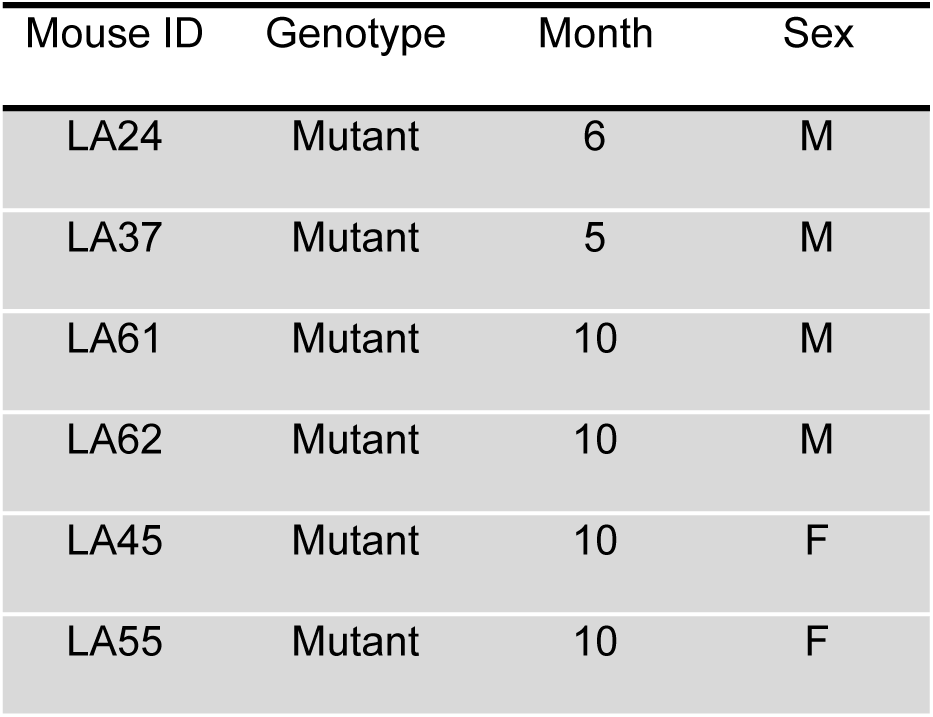
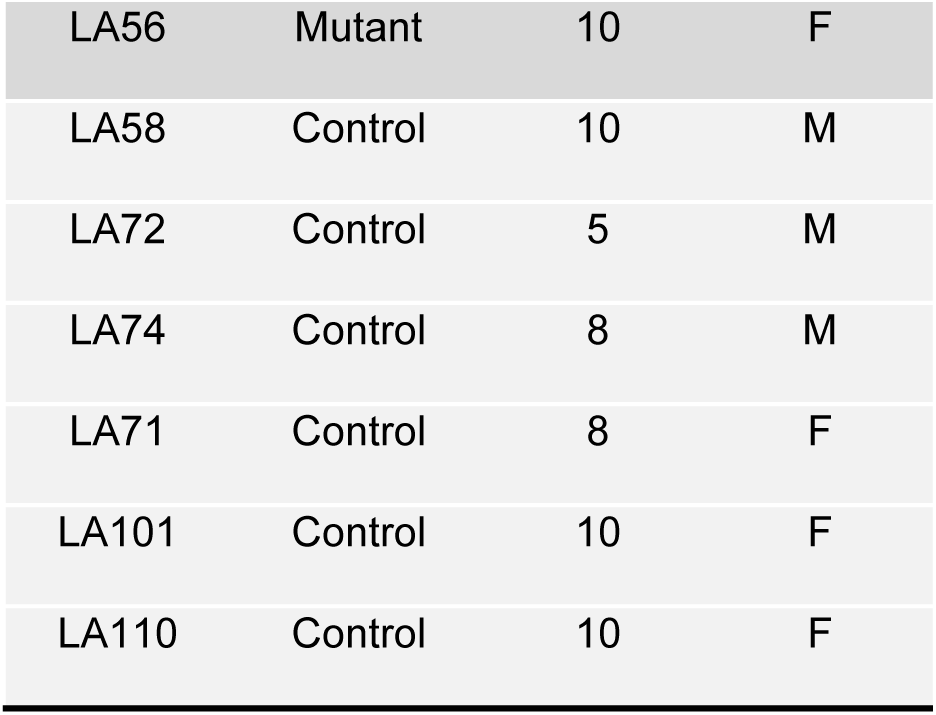
Mouse model of tauopathy and controls used for functional analyses.

**Table 2.**
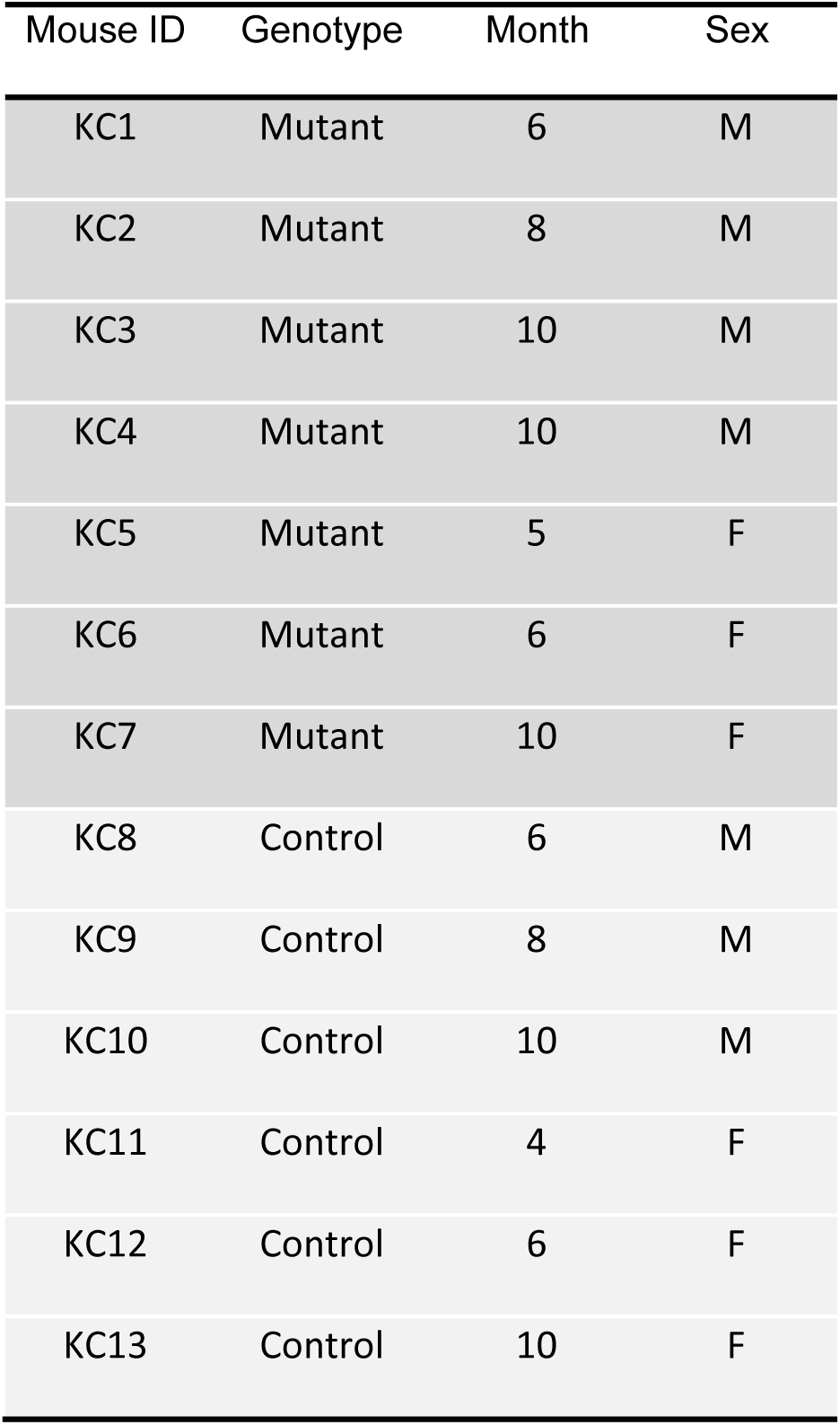
Mouse model of tauopathy and controls used for immunohistochemical analyses.

### METHOD DETAILS

#### Surgery

A 4-mm diameter craniotomy was performed over visual cortex as previously described^67^. Briefly, mice were first sedated with acepromazine (2 mg/kg body weight, i.p.) and then were deeply anesthetized using isoflurane (2% for induction, 1-1.5% for surgery). The body temperature was monitored and actively maintained using an electronic heat pad regulated via rectal probe. Carprofen (20 mg/kg body weight, s.c.) was administered preoperatively, and lidocaine solution containing epinephrine (5 mg/kg body weight s.c.) was injected locally before and after the scalp excision. The scalp overlaying the right visual cortex was removed and a custom head-fixing imaging chamber with a 5-mm diameter opening was mounted to the skull with cyanoacrylate-based glue (Oasis Medical) and dental cement (C&B Metabond, Parkell). Mice were mounted on a custom holder via the headplate chamber, which was filled with a physiological saline containing (in mM) 150 NaCl, 2.5 KCl, 10 HEPES, 2 CaCl2 and 1 MgCl2. A craniotomy was performed using a carbide dental bur on a contra-angle handpiece (NSK).

#### Intrinsic signal optical imaging (ISOI) and retinotopic maps

In order to functionally map visual cortex for targeted injection of viral vectors, ISOI was performed using a custom macroscope and a CCD camera as previously described^67,68^. Briefly, the macroscope was focused down 600 μm from the pial surface, and intrinsic signals were illuminated with halogen light (Asahi Spectra) delivered via light guides and focusing probes (Oriel) through a red filter (700 ± 38 nm; Chroma). Reflected light was then captured through a second red filter (700 ± 5 nm; Edmund Optics) at a 30Hz frame rate with a custom image acquisition software (original code kindly provided by D. Ferster, Northwestern University and adapted in Smith et al., 2017). Intrinsic signals were quantified as a change in the captured reflected light at 700 nm that correlated with the cyclic pattern of the visual stimuli. During imaging, mice were lightly anesthetized with 0.25 – 0.5% isoflurane augmented by acepromazine (2 mg/kg body weight, i.p.), and the body temperature was maintained at 37 °C. A single drifting white bar on a black background (3° thick, moving in elevation or azimuth direction) was used as previously described^67^ in order to obtain retinotopic maps.

#### Viral injections

Retinotopic maps determined via ISOI were used to locate V1. The pial vasculature map relative to the retinotopic maps was used to guide targeted injections into V1. A combination of two adeno-associated viral (AAV) vectors encoding Cre-recombinase and flexed genetically encoded calcium indicator (GECI), GCaMP8m, respectively, were injected into V1 under continued surgical plane of isoflurane anesthesia44-46. Briefly, 1:1 mixture of pENN.AAV.CamKII 0.4.Cre.SV40 (AAV1; Addgene #105558; diluted at 1:20,000 in phosphate buffered saline (PBS) with final concentration at 5 x 108 vg/ml) and pGP.AAV.syn.FLEX.jGCaMP8m.WPRE (AAV1; Addgene #162378; original concentration at ∼1013 vg/mL) viral particles were injected (65 - 100nL per site; 1∼2 sites per animal) with a pulled-glass capillary micropipette using a Nanoliter 2010 controlled by a microprocessor, Micro4 (World Precision Instruments), at 15 nL per min. The glass pipette was left in place for 5 mins before retracting to avoid the backflushing of the injected viral solution. The cranial window was then sealed with a glass cranial plug made up of 4-mm and 3-mm circular coverslips (Warner Instruments) stacked in tandem with a UV-curing optical adhesive (NOA61, Norland).

#### Visual stimuli for assessing orientation tuning

A small LCD monitor placed 10cm away from the animal was used to deliver visual stimuli during 2p-Ca2+ imaging. In order to prevent the stimulus light from contaminating the imaging pathways, the visual stimulus LCD monitor was covered with a custom 3D-printed pyramid-shaped shroud with a small opening at the tip that was positioned around the eye of the mouse. The stimulus extended from +20° to +124° in azimuth and from-10° to +42° in elevation. Stimulus frames consisted of 128 x 128 pixels, which were smoothly interpolated such that one pixel was equivalent to 0.72°2 in visual space. Visual stimuli were generated using MATLAB and the Psychophysics toolbox^69^. Orientation tuning was assessed using drifting black and white square wave gratings as previously described^45^ (12 different directions with 30° increments or 8 different directions with 45° increments, at 0.04 cycles/° and 2Hz). Each grating was presented in a random order for 3 s separated by 1 s of gray screen in between for each sweep, which was then repeated 8 times. Two different contrast levels (40 or 100%) were used for each neuron that would elicit enough spine signals while reducing the invasion of backpropagating action potential (bAP) into the spines. Therefore, the response property measurements were normalized to the average of contrast-matched controls. Neuronal output was considered visually responsive if the following cut-off criteria were met: mean peak angle activity ≥ 0.2 dF/F and all stim responses had >1 signal to noise ratio compared to all responses to the latter half (0.5s) of the grey screen presentation. Because the spine-specific responses that remain after bAP signal removal was small and the frequency and the amplitude of bAP to be removed was variable, spine response was z-scored to normalize their activity to the baseline fluctuations. Dendritic spine activity was considered visually responsive if the following cut-off criteria were met: mean peak angle activity ≥ 0.5 z-score and all stim responses had > 1 signal to noise ratio compared to all grey screen responses. For both dendritic and spine, visual responses were first fit to a sum of two Gaussian curves with the two peaks set 180° apart. Dendrites and spines were considered orientation tuned if a sum of two Gaussian curves with the two peaks set 180° apart fit successfully with r^2^ ≥ 0.7 (**Figure S6).** The curve containing the preferred orientation was used to produce orientation tuning curves.

#### Two-photon microscopy imaging

2p-GCaMP8m imaging was performed starting 4-6 weeks after AAV injection, using a custom-built 2p-microscope used in prior studies^43,45,67^. In order to achieve the stability crucially required for dendritic imaging with an individual spine resolution, animals were lightly anesthetized with 0.25 to 0.5% isoflurane augmented with 1mg/kg acepromazine or with 0.5 to 1.0% isoflurane without acepromazine. Frame scans were acquired with a 16x magnification, 0.8 numerical aperture water immersion objective (Nikon) using ScanImage^70^ at 30 Hz (512×512 pixels), 58 Hz (512×256 pixels), or 110 Hz (512 x 128 pixels). The total number of frames acquired per each imaging session was 13500, 25500, and 42000 frames for 30°-increment gratings and 9000, 17000, and 33000 frames for 45°-increment gratings at 30, 58, and 110Hz scanning, respectively. Average excitation laser power measured at the objective was 65.1 ± 3.7 mW. For dendritic activity, we used CASCADE pretrained model for GCaMP8m^71,72^ to calculate inferred total spike count during the visual stimulation period.

#### Structural characterization of dendritic spines

To determine dendritic spine density and spine turnover, GCaMP8m-expressing dendrites were imaged on two timepoints separated by 3-16 days. The average interval was not statistically different between the control and mutants (control 8.79 ± 0.69 vs. mutant 7.93 ± 1.24, two-tailed Wilcoxon p = 0.4689). A custom dendritic spine analysis software, AUTOTUNE, was used for in-plane motion correction, region of interest (ROI) segmentation, and extraction of GCaMP8m transients. The spine turnover module of the AUTOTUNE was used to analyze the spine positions along the dendrite, perform one-to-one comparison of the imaged spine locations across the two timepoints and designate them as retained, gained, or lost spines^45^ (https://github.com/yuyiyi/AUTOTUNE_GUIdevelopment.git). The outputs from the AUTOTUNE were also visually inspected and corrected, if necessary, by trained experts. The spine elimination rate was calculated as the number of lost spines divided by the total number of spines present in the first timepoint (lost spines/(retained + lost spines)) for each neuron. Similarly, the spine formation rate was calculated as the number of gained spines divided by the total number of spines present in the first timepoint (gained spines/(retained + lost spines)). Spine turnover ratio was calculated as the average of elimination and formation rate for each neuron ((lost + gained spines)/(retained + lost spines)*2). Additionally, in order to normalize the potential impact of varying intervals between the two imaging sessions, the turnover ratio per day was also calculated (turnover ratio/timepoint interval (days)). Further inspection of the spine ROIs that turned over revealed functional properties of those spines that met the specific fate compared to others.

#### Functional characterization of synaptic inputs

Fluorescent transients were measured as the mean of all pixels within the ROI at each frame. For functional characterization of dendritic spines, the input mapping module of the AUTOTUNE was used to remove the signal contamination by the backpropagating action potentials^45^ (bAPs). Briefly, the module adopted the method by Chen et al., 2013^73^ in which a parent dendritic shaft signal is linearly subtracted from the spine signals at a scale set for each and individual spine. The extracted true spine-specific activity was analyzed in order to determine the orientation tuning property of the individual spines^45^. The orientation selectivity index (OSI) was computed using the tuning curves generated based on the fluorescent time-course:

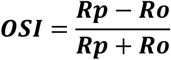

where Rp and Ro are the responses to the preferred and orthogonal orientations, respectively. Preferred orientation, tuning width (FWHM), and Δ preferred orientation relative to the tuning of the parental neuronal activity were determined after fitting a sum of two Gaussian curves, with the respective peaks set to be 180° apart, to the tuning curves. The curve containing the preferred orientation was used to produce orientation tuning curves. OSI and FWHM values were then normalized to the average of visual-stim contrast-matched controls.

#### Immunohistochemistry

Mice were euthanized with a mixture of ketamine (300 mg/kg), xylazine (60 mg/kg), and azepromazine (9mg/kg) and transcardially perfused with cold PBS followed by 4% paraformaldehyde (PFA) solution. Brains were dissected out, postfixed overnight in 4% PFA, and cryoprotected in a series of sucrose solutions with increasing concentration (until tissue sank in 15% and subsequently in 30% sucrose in PBS) before being stored at-80 °C.

Coronal slices (40µm thick) were prepared using a Leica CM1860 UV cryostat (Leica Biosystems) processed for immunohistochemistry with the following antibodies as previously described^74^ with slight modifications as described below. Primary: chicken polyclonal anti-MAP2 (Abcam, ab5392) and mouse monoclonal IgG1 anti-phospho Tau (AT8; ThermoFisher Scientific, MN1020) antibodies; Secondary: Alexa Fluor 488 goat anti-chicken IgY (H+L) antibody (Invitrogen, A-11039), and Alexa Fluor 594 goat anti-mouse IgG (H+L) antibody (Invitrogen, A-11005). Briefly, slices were washed in PBS (3x 10 minutes), and blocked with 10% Bovine serum albumin (BSA; Millipore Sigma, A3294), 0.5% triton X-100 (Millipore Sigma, 93443), and PBS for 1 hour at room temperature. Primary and secondary antibodies were diluted in 10% BSA and 0.5% triton X-100 in PBS. Slices were incubated with the primary antibodies (1:1000 dilution) for 48 hours at 4°C after which were washed in PBS (3x 10 minutes) and incubated in the secondary antibodies (1:500 dilution) for 1.5-2 hours at room temperature. Slices were then washed in PBS (3x 10 minutes), mounted onto a slide, and coverslipped with mounting medium with DAPI (Vector Laboratories, H-1500-10).

#### One-photon microscopy

Slice images were captured using a Keyence BZ-X1000 Fluorescence Microscope (4x and 10x objective), a Leica SP8 Resonant Scanning Confocal Microscope (20x and 40x objective), and an Abberior Instruments STED Microscope (60x objective). Images taken with the 4x objective were stitched together using Keyence stitching software while others taken with the 20x and 40x objectives were stitched together using Leica stitching software. Allen Reference Atlas (Mouse Brain)^75^ was used as anatomical reference to locate V1.

#### Immunohistochemistry quantification

Images were aligned so that the surface of the brain was horizontal. To measure layer 2/3 phosphorylated tau expression, a 150 µm by 150 µm square region of interest (ROI) was selected manually in layer 2/3 of V1, which was identified with DAPI staining. The average intensity of AT8 immunostaining in the ROI was measured using ImageJ^76^. To measure the profile of phosphorylated tau expression along the cortical depth, a 100 µm by 800 µm rectangular ROI was selected in V1 perpendicular to the pial surface, spanning the cortical layers. The top of the ROI was aligned at the surface of the cortex. The profile of AT8 intensity along cortical depth was measured by averaging the raw fluorescence along the tangential dimension of the cortex in the ROI. For plotting the intensity along cortical layers, layer boundaries were defined according to previously reported depths^77^. 2 slices were analyzed per mouse for all immunohistochemistry quantification.

#### Classifier models and evaluation

Classifiers were trained on five dendrometric parameters, global bAP-inferred spiking activity in the dendrites, AUC, OSI, FWHM, and spine density. 10-fold cross validation was used to avoid overfitting, and the trained models were tested on the held-out 15% of data. We trained and tested the performance of all 34 models available on the MATLAB Classification Learner App over 8 trials each and selected three models, Gaussian Naïve Bayesian, Quadratic Discriminant, Quadratic Support Vector Machine (SVM), for their reliable validation and test accuracy. To evaluate the performance of these trained models, Receiver Operating Characteristic (ROC) curves were plotted, and the Area Under the Curve (AUC) was computed. In order to determine the impact of each predictor parameter, permutation feature importance (PFI) was calculated. 10 permutations were performed to ensure a robust estimate, and the mean importance and standard error were plotted.

### QUANTIFICATION AND STATISTICAL ANALYSIS

Unless otherwise stated, all measurements in the text are presented as means ± s.e.m., and violin plots are presented with a white dot indicating the mean. The Shapiro–Wilk test was used, before the application of parametric tests, to confirm that the deviation of the data from normality was statistically insignificant. For non-normal data, or data with significantly different variances, non-parametric tests were used as noted in the text. For multiple group comparisons of non-normal data, Dunn’s multiple comparison test was used, controlling for Type I errors. Sample sizes were designed to reliably measure neurophysiological parameters while conforming to ethical guidelines and minimizing animal use. Key statistical details of the experiments can be found on Table 3 and in result sections. No randomization was used in analyses. For all analyses, the experimenters were blind to the genotype or the identity of the animals from which the data were collected.

**Table 3.**
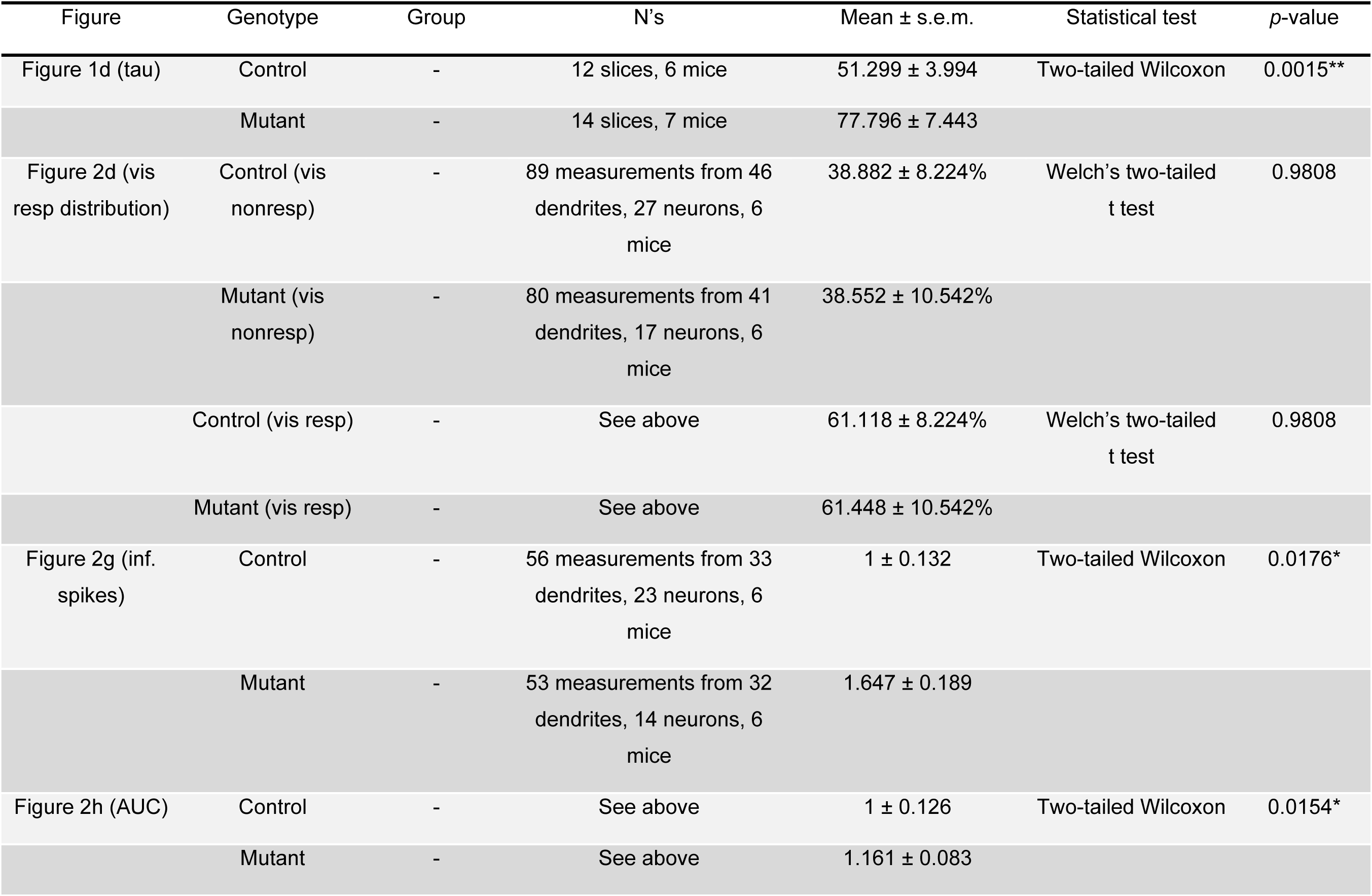
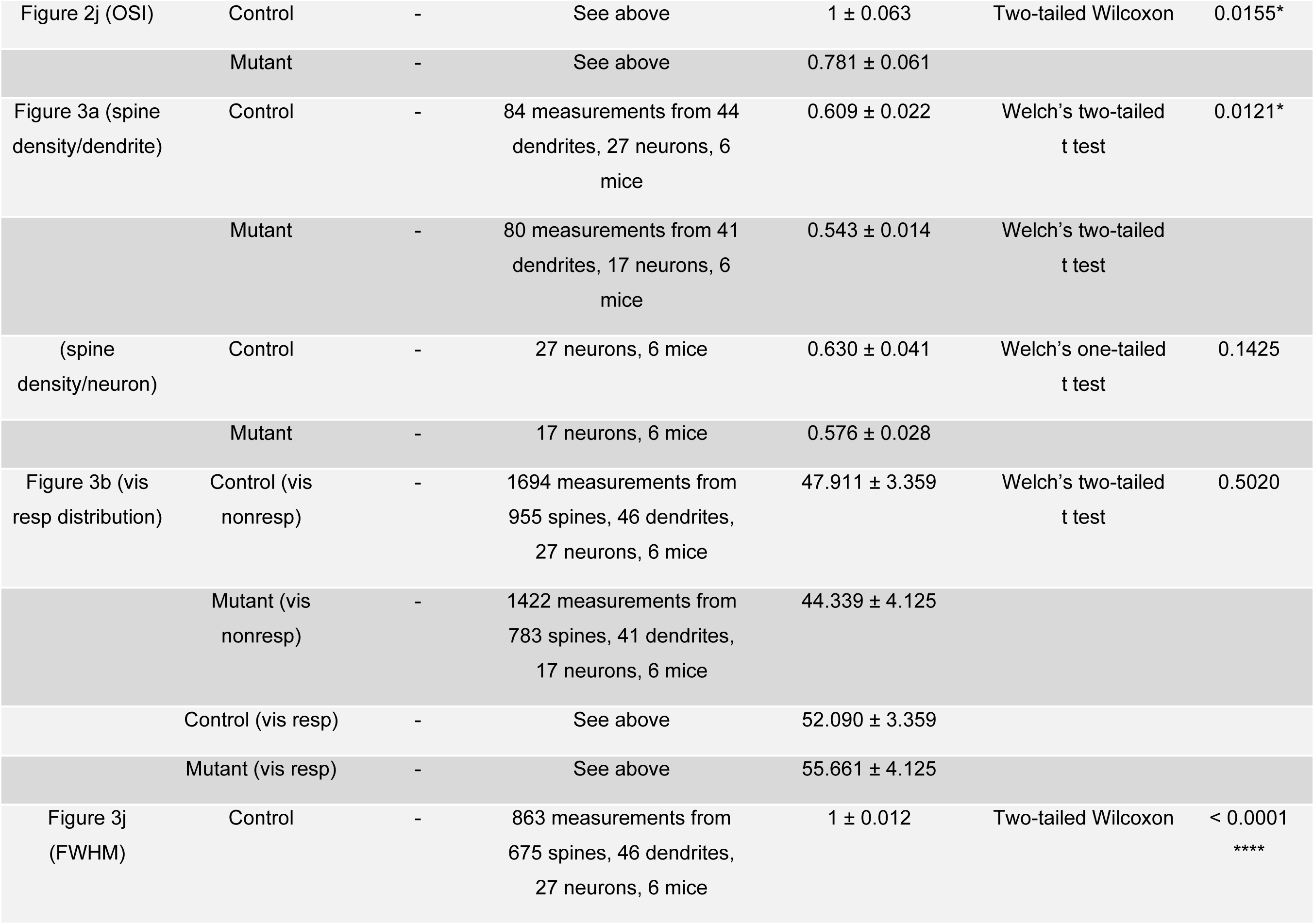
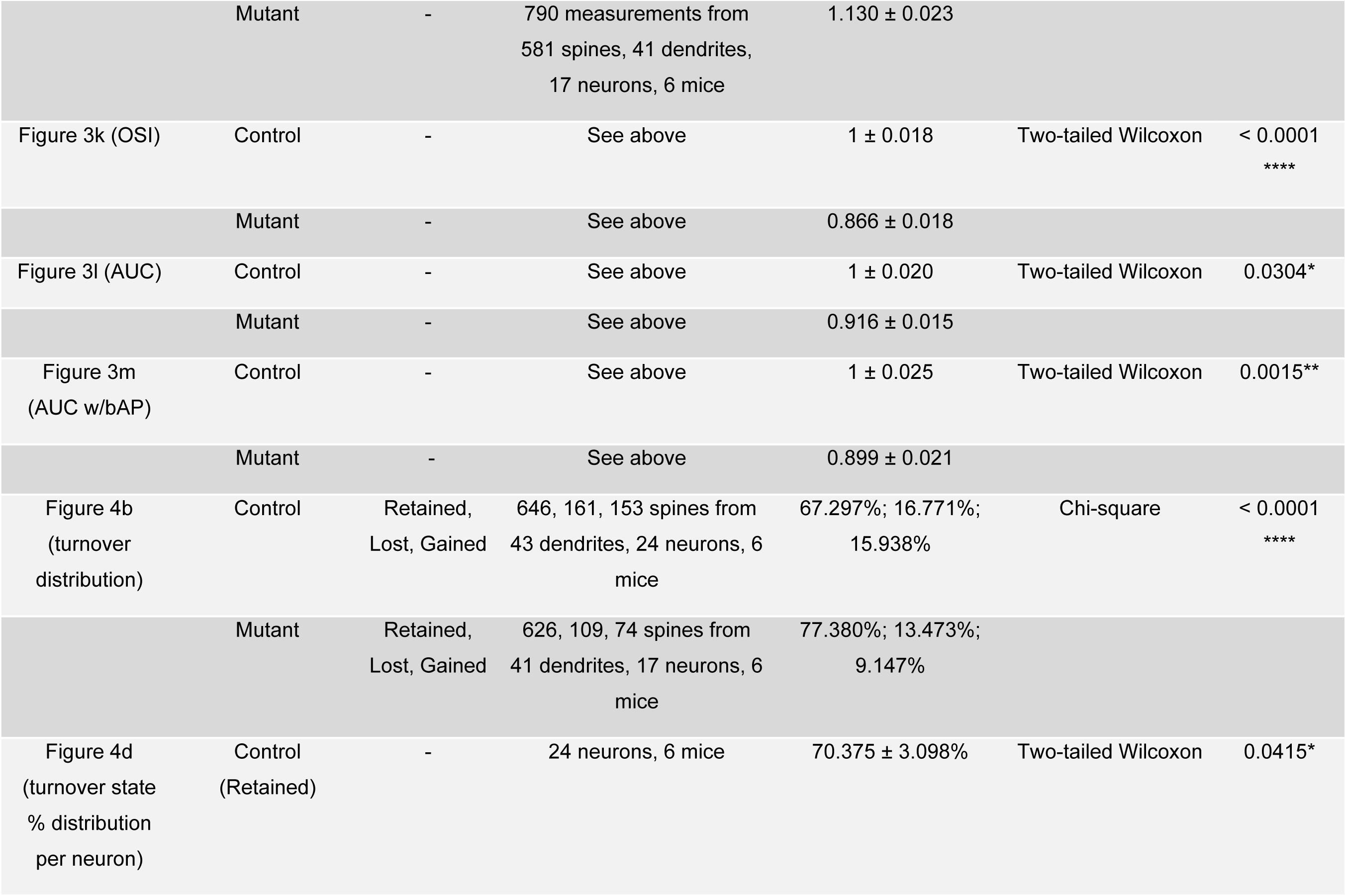
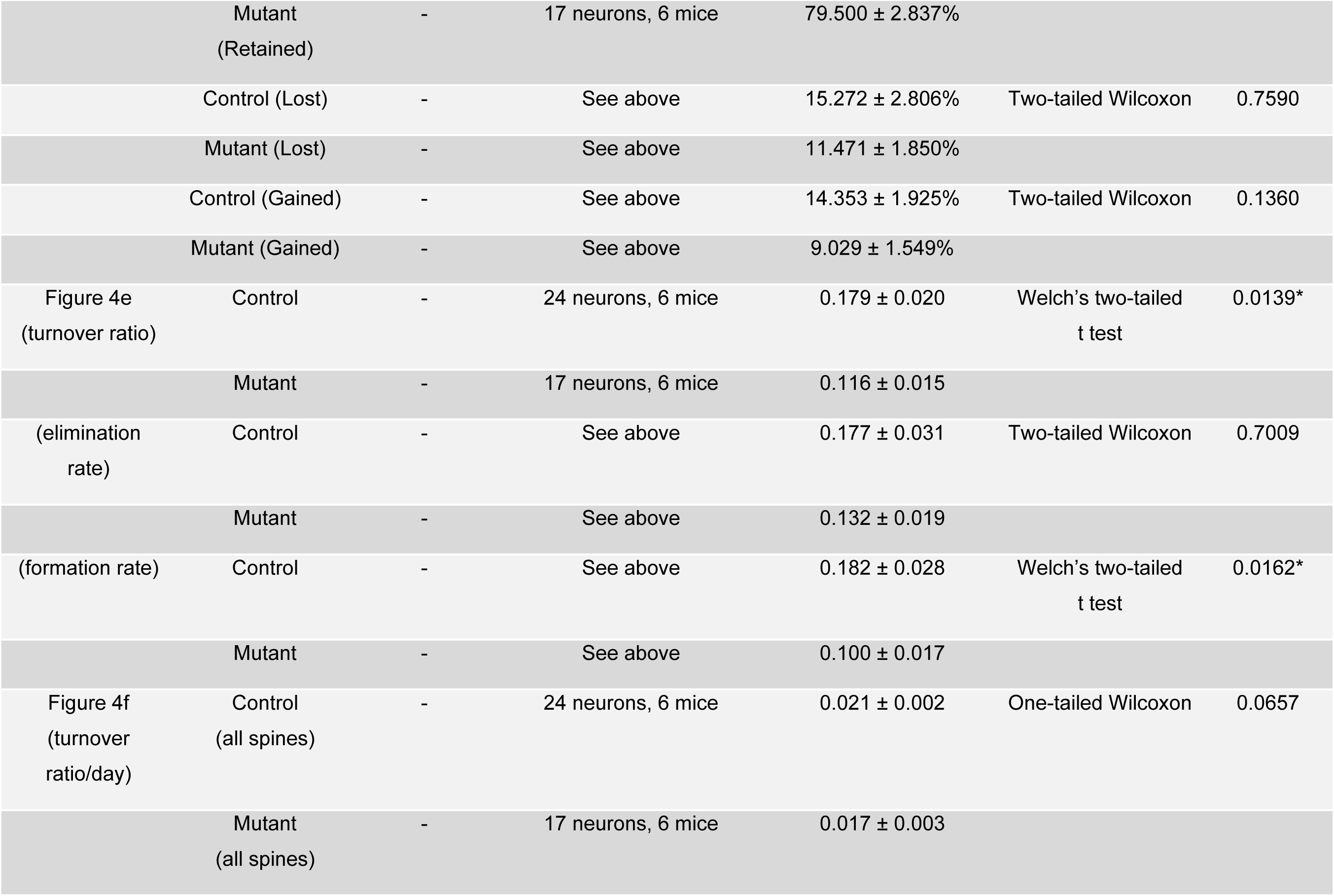
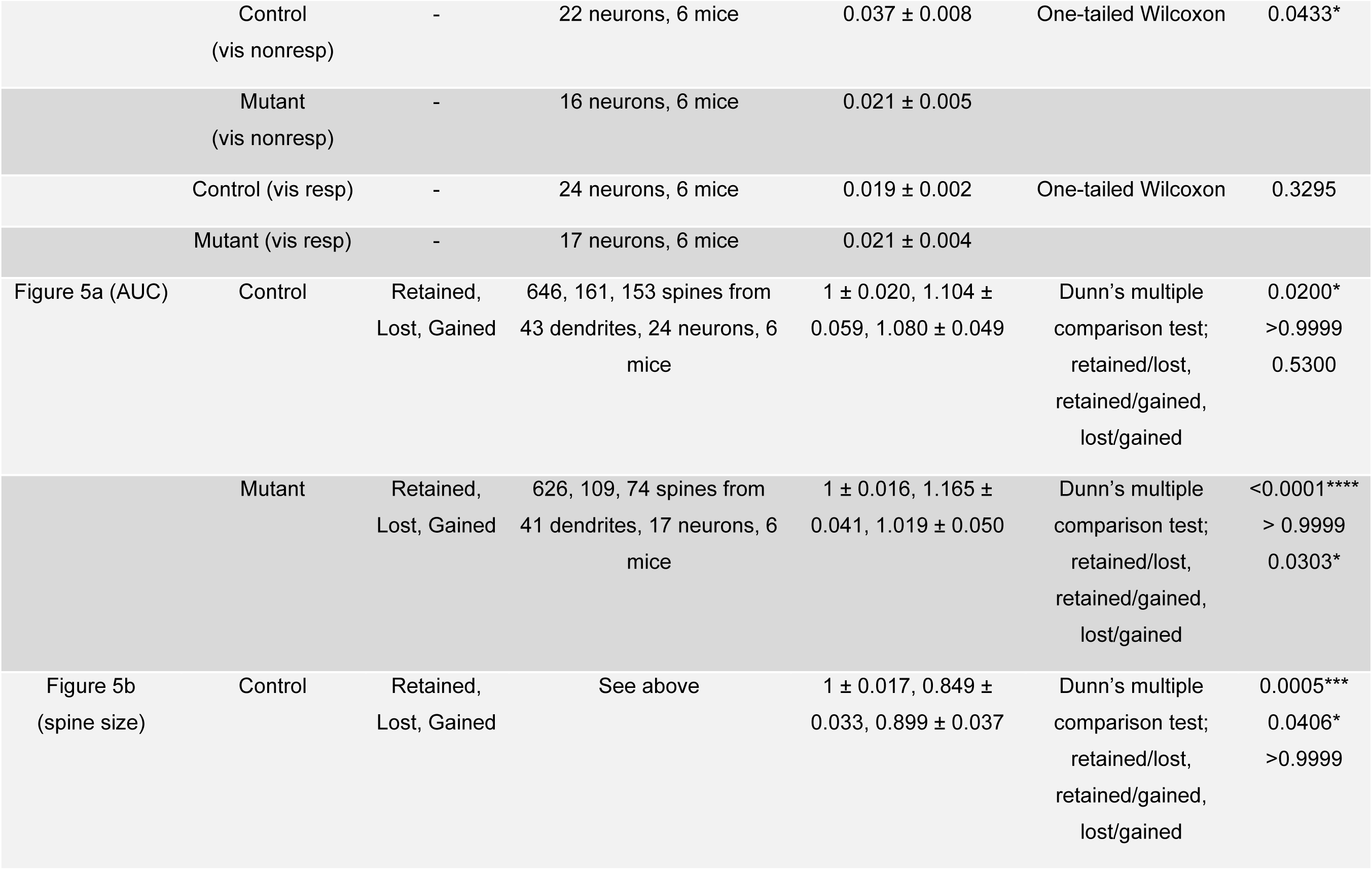
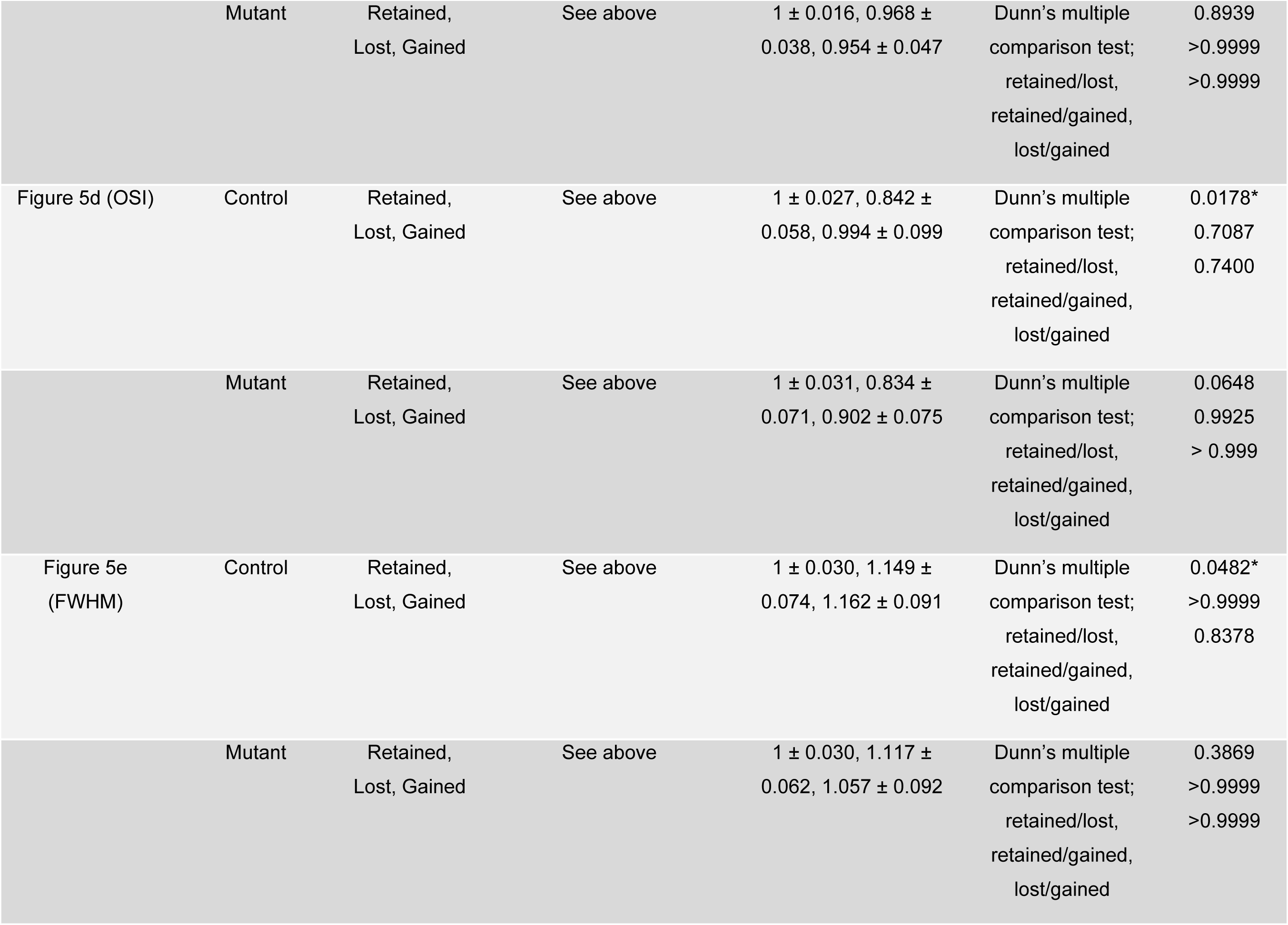
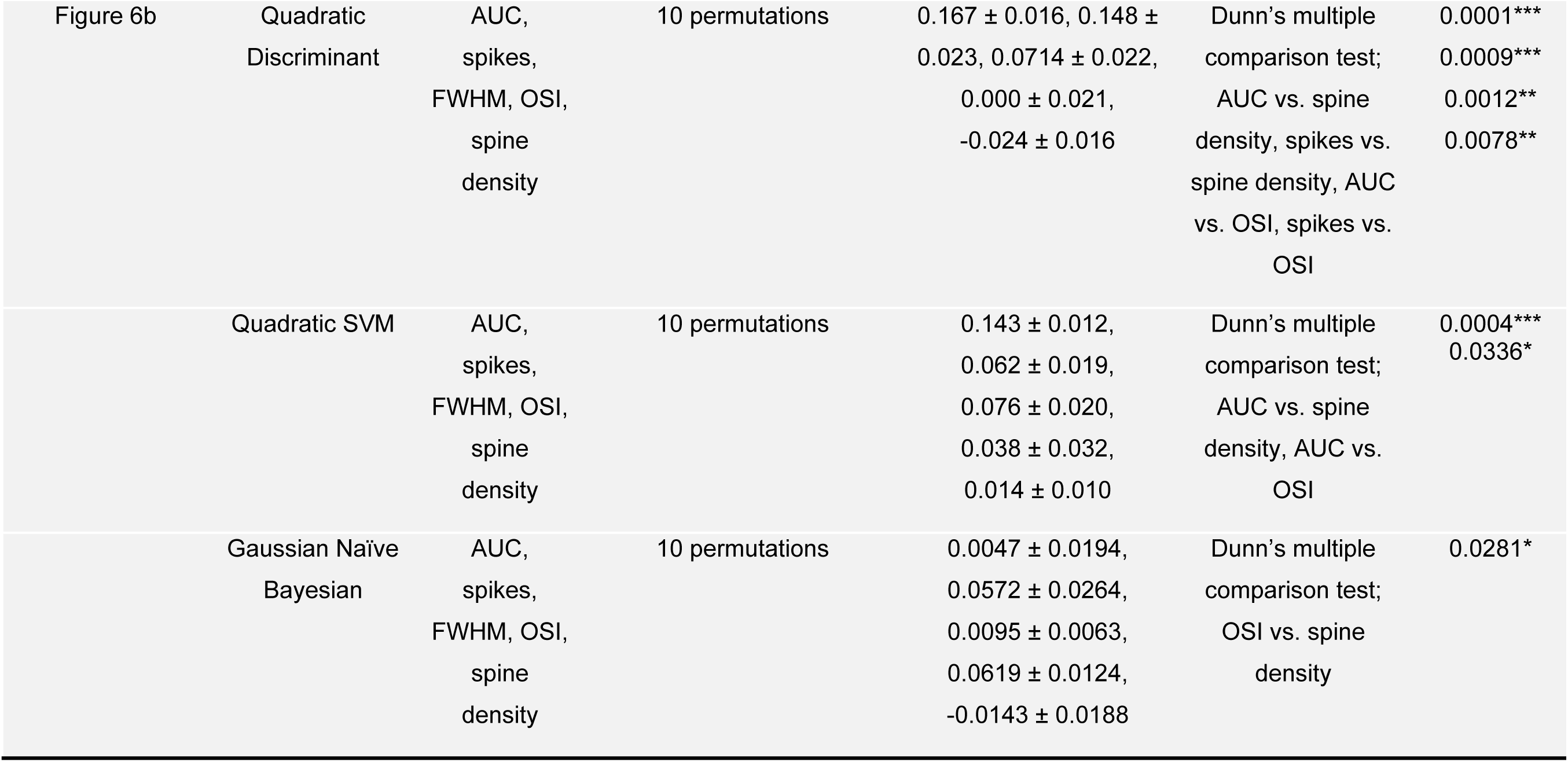
Key statistics.

---

**Figure S1.**
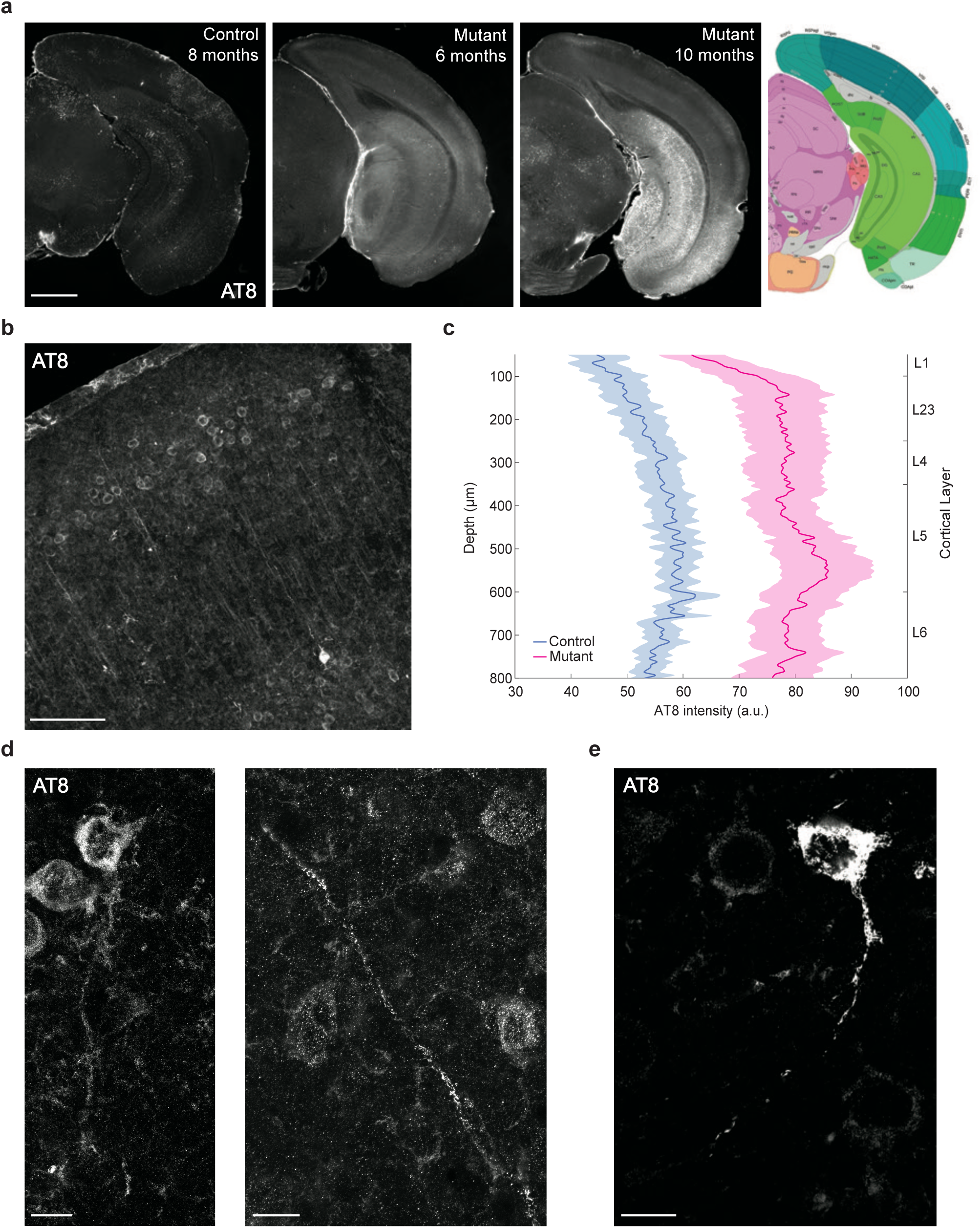
Pathological tau expression in P301S mutants. (A) Left: Example coronal slices show robust expression of phosphorylated tau labeled with AT8 antibodies in P301S mutant mice (6 and 10 months) in the hippocampal formation (e.g. dentate gyrus, hippocampus, entorhinal cortex) and more modest expression in the neocortex that are not present in controls (8 months). A slice from the Allen Mouse Brain Atlas (ref.75) corresponding to equivalent coordinates shown for an anatomical reference. Scale bar = 1 mm. (B) Phosphorylated tau expression across the cortical layers of V1 in mutant mouse (6 months), showing robust expression in somas and in apical dendrites of layer 5 neurons. Scale bar = 100 µm. (C) Quantification of V1 phosphorylated tau expression by cortical depth for control (n = 12 slices) and mutant mice (n = 14 slices). (D) Example dendrites with phosphorylated tau expression (6-month mutant). Scale bar = 10 µm. (E) Example dendrite with phosphorylated tau expression (10-month mutant). Scale bar = 10 µm.

**Figure S2.**
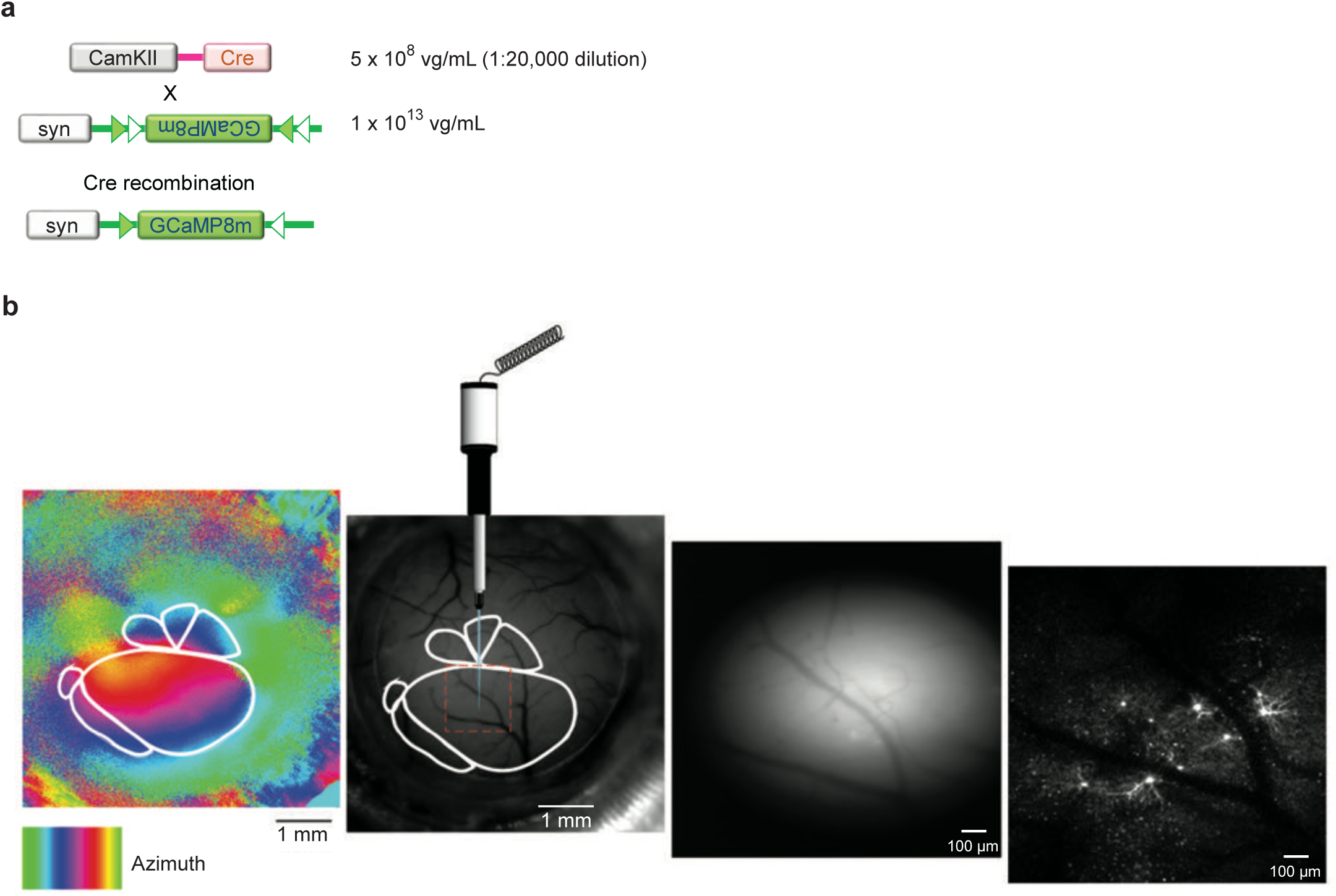
Functionally targeted injection of AAV vectors for ultra-sparse transduction in V1. **a.** A combination of two AAV vectors encoding Cre-recombinase and flexed GCaMP8m were used. AAV-Cre was diluted by x 20,000 to achieve sparse expression. **b.** Retinotopic maps obtained with ISOI were used to target V1 for AAV injection. Large cortical vessels served as a landmark to locate the site of injection and the transfected neurons expressing GCaMP8m.

**Figure S3.**
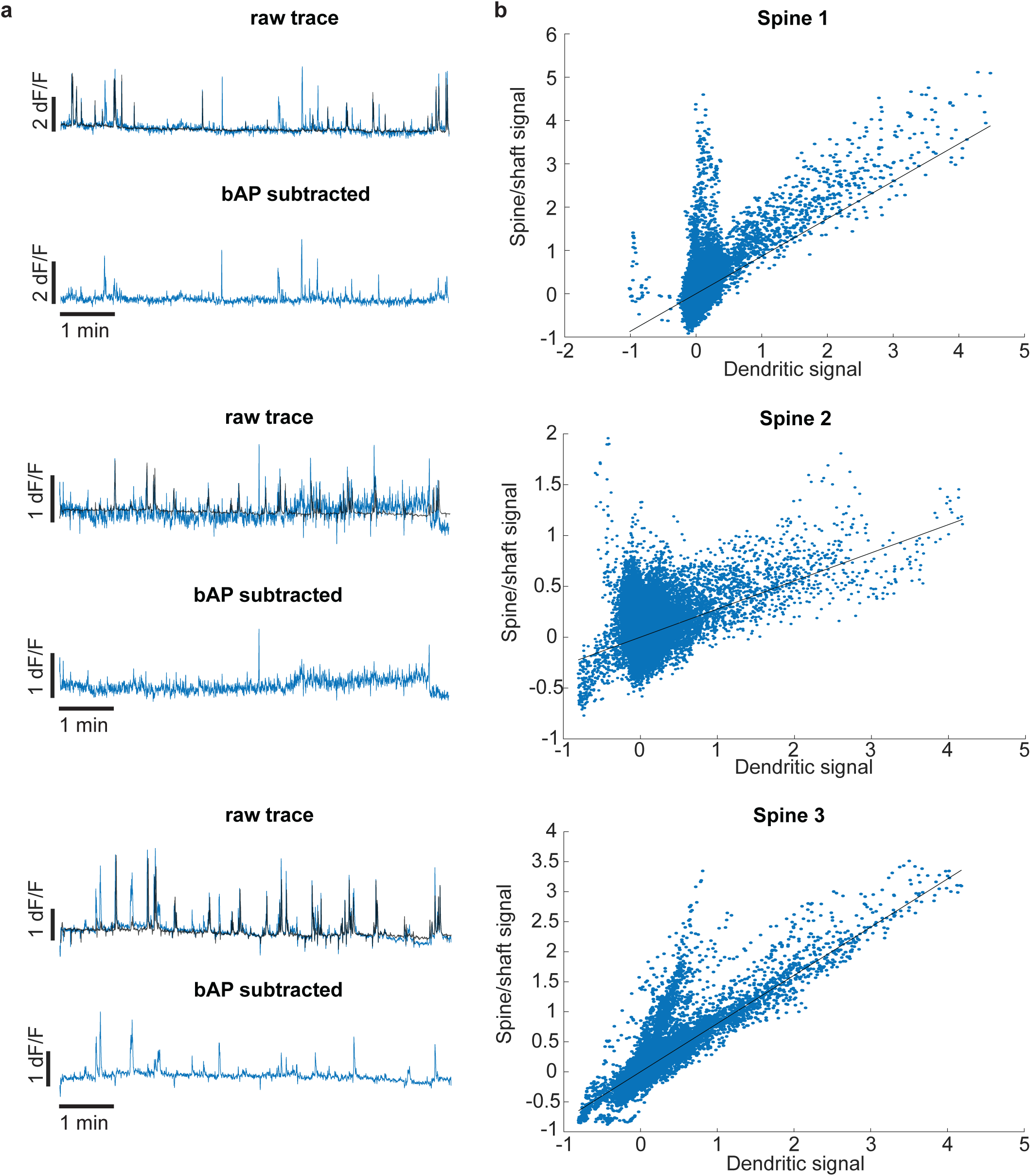
bAP subtraction for isolating dendritic spine-specific signals. **a.** Three example sets of calcium transient traces for spines 1 - 3 (blue traces) overlaid with concurrent bAP signals measured from the parent dendrite (black). bAP signals were scaled down by the factor calculated in **b** and subtracted from the raw spine traces to isolate spine-specific transients (bAP subtracted). **b.** Using AUTOTUNE, robust fit function was applied to the correlated portion of the spine/dendritic activity to calculate the appropriate scale factor (slope).

**Figure S4.**
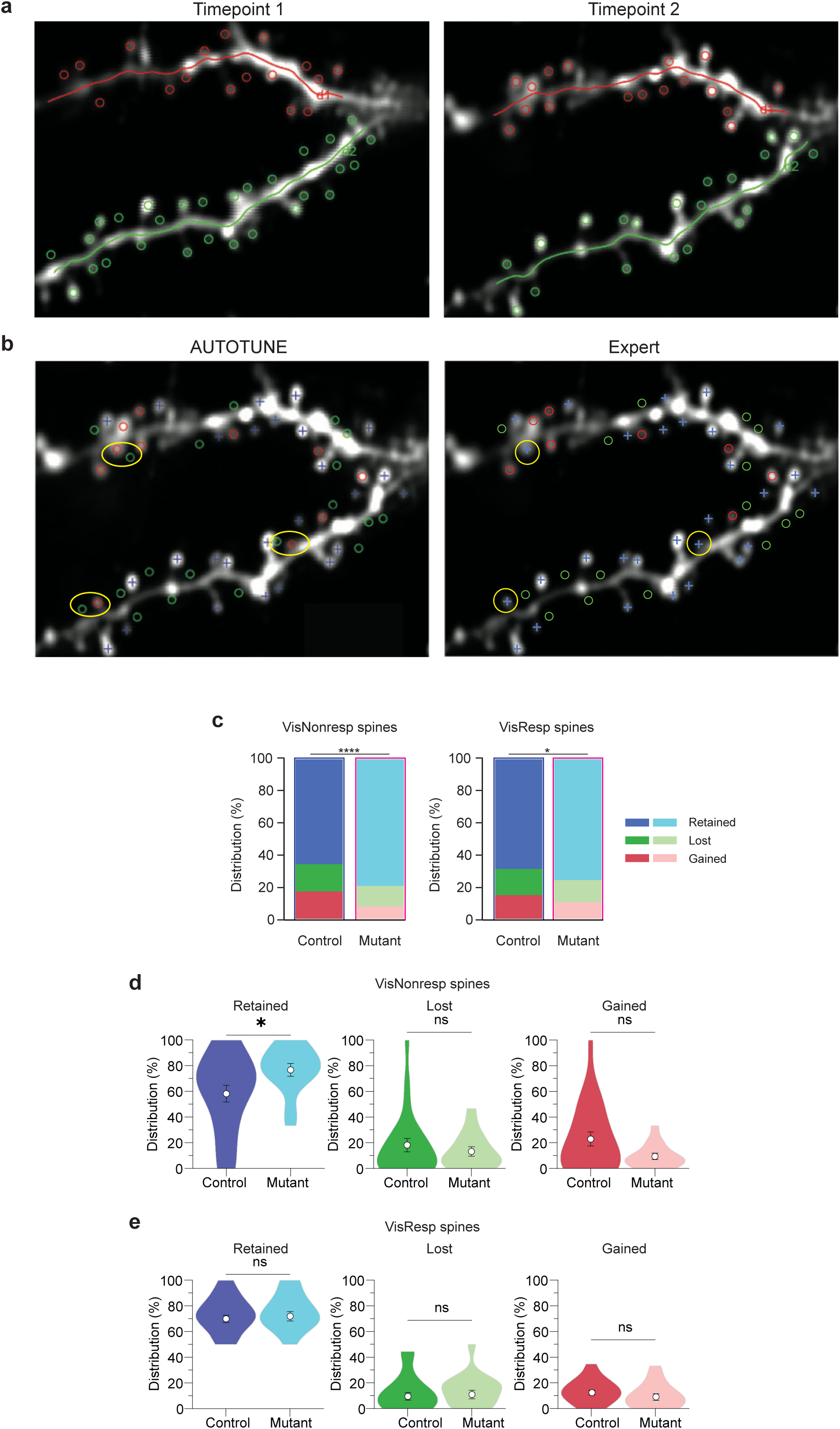
Automated vs. manual assessment of spine turnover. **a.** Regions of interests (ROIs) drawn for dendrite 1 (d1, red) and dendrite 2 (d2, green) and their spines at timepoint 1 (left) and timepoint 2 (right), separated by 7 days. **b.** Designation of spines as retained (blue cross), lost (green circle), or gained (red circle) performed by the automated software AUTOTUNE (left) and human expert (right). Yellow circles highlight the mismatch in the designation where human expert was superior in identifying a retained spine while the automation defined them as a combination of lost and gained spines in close proximity. **c.** The distributions of retained, lost, and gained spines were significantly different between controls and mutants, for visually non-responsive and responsive spines. **d.** The percentage of retained visually non-responsive spines per neuron in mutant was significantly higher than that of controls. **e.** Visually responsive spines have consistent proportions for retained, lost, and gained across genotype.

**Figure S5.**
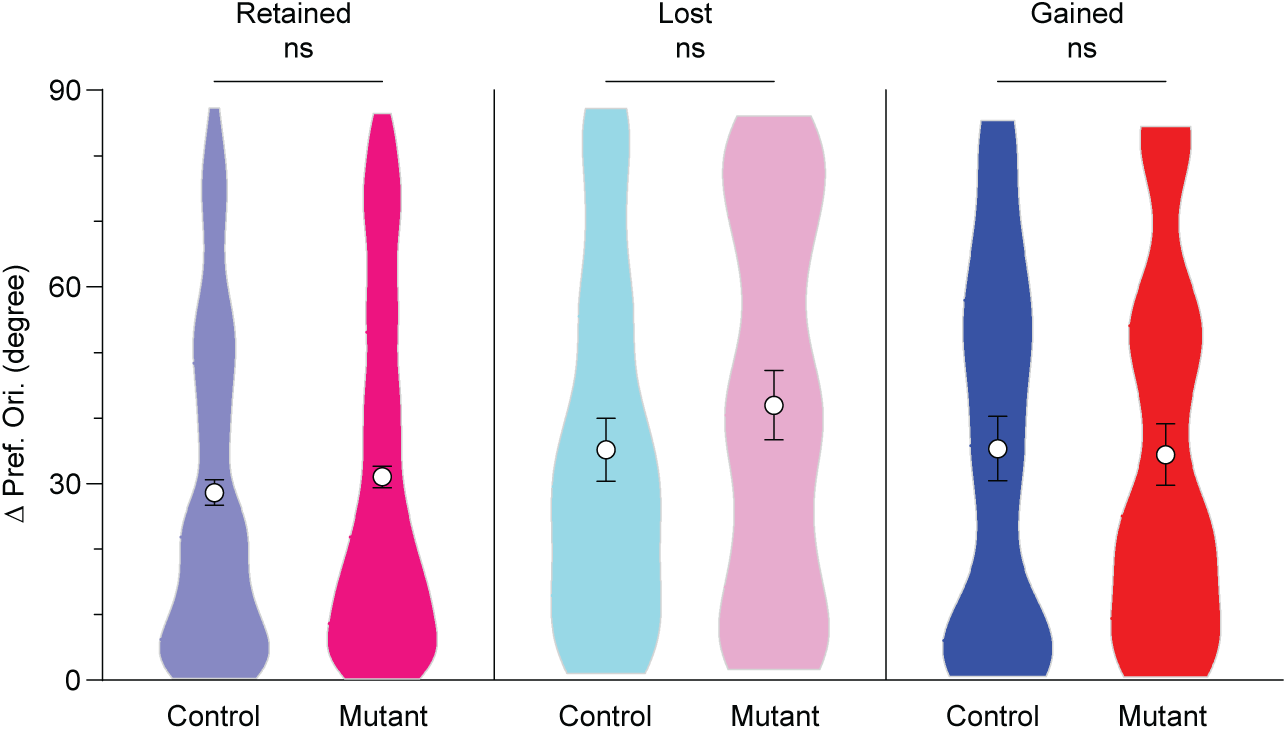
Distribution of preferred orientations for retained, lost, and gained compared to neuronal output. Average preferred orientations were similar between spines in mutant and control mice regardless of the turnover status.

**Figure S6.**
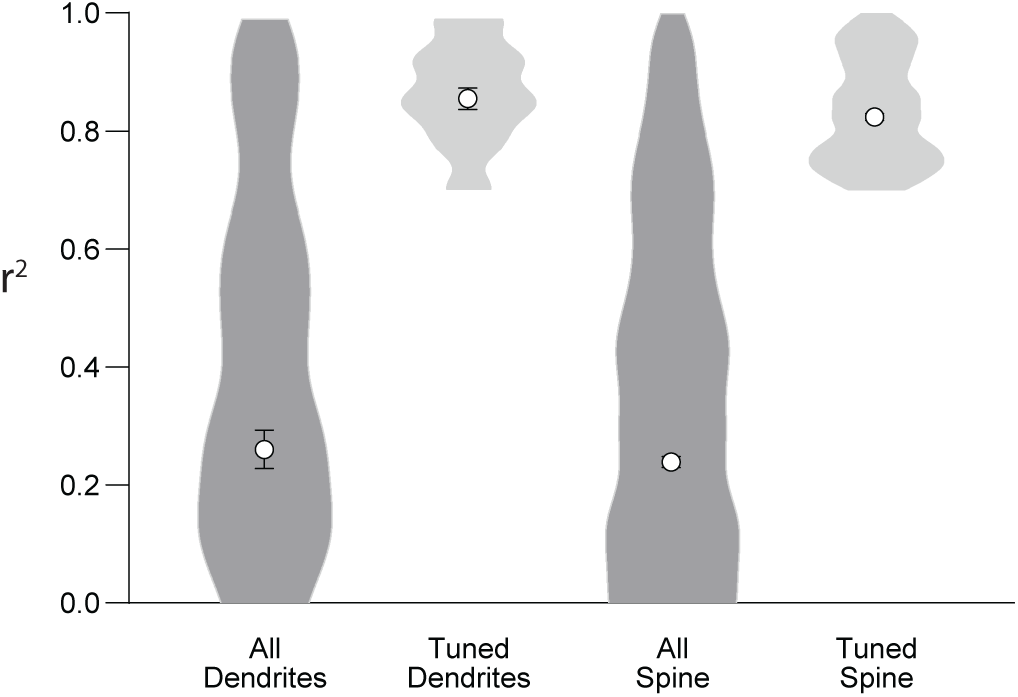
Coefficient of determination (r²) calculated as a measure of Gaussian fit. Those dendrites and spines whose fit exhibited r^2^ 2: 0.7 were considered visually tuned.

